# A PI3Kδ-Foxo1-FasL signaling amplification loop rewires CD4^+^ T helper cell signaling, differentiation and epigenetic remodeling

**DOI:** 10.1101/2024.10.28.620691

**Authors:** Dominic P Golec, Pedro Gazzinelli-Guimaraes, Daniel Chauss, Hiroyuki Nagashima, Kang Yu, Tom Hill, Luis Nivelo, Jennifer L Cannons, Jillian Perry, Ilin Joshi, Nicolas Pereira, Fabrício Marcus Silva Oliveira, Anthony C Cruz, Kirk M Druey, Justin B Lack, Thomas B Nutman, Alejandro V Villarino, John J O’Shea, Behdad Afzali, Pamela L Schwartzberg

## Abstract

While inputs regulating CD4^+^ T helper cell (Th) differentiation are well-defined, the integration of downstream signaling with transcriptional and epigenetic programs that define Th-lineage identity remain unresolved. PI3K signaling is a critical regulator of T cell function; activating mutations affecting PI3Kδ result in an immunodeficiency with multiple T cell defects. Using mice expressing activated-PI3Kδ, we found aberrant expression of proinflammatory Th1-signature genes under Th2-inducing conditions, both *in vivo* and *in vitro*. This dysregulation was driven by a robust PI3Kδ-IL-2-Foxo1 signaling loop, fueling Foxo1-inactivation, loss of Th2-lineage restriction, altered chromatin accessibility and global impairment of CTCF-DNA interactions. Surprisingly, ablation of *Fasl*, a Foxo1-repressed gene, restored normal Th2 differentiation, TCR signaling and CTCF expression. BioID revealed Fas interactions with TCR-signaling components, which were supported by Fas-mediated potentiation of TCR signaling. Our results highlight Fas-FasL signaling as a critical intermediate in phenotypes driven by activated-PI3Kδ, thereby linking two key pathways of immune dysregulation.

## Introduction

The differentiation of CD4^+^ T cells into distinct types of cytokine-producing T helper (Th) cell populations is critical for orchestrating immune responses to different types of infections and immune challenges. Among these, Th1 cells express IFNγ, which drives activation of cellular immunity against intracellular pathogens, whereas Th2 cells produce IL-4, IL-5 and IL-13, which promote both hypersensitivity and tissue repair processes. T helper cell differentiation requires the translation of extracellular signals from the T cell receptor (TCR) and cytokines into intracellular responses that dictate transcriptional and epigenetic signatures required to establish T cell identity. Nonetheless, how these pathways coalesce to regulate CD4^+^ Th cell fate remains incompletely understood.

Phosphoinositide 3-Kinase (PI3K), is a major cellular signaling hub that helps orchestrate T cell responses downstream of the TCR, cytokine receptors and chemokine receptors. Coordination and amplification of T cell signaling, metabolism and transcription are key features of PI3K. The importance of PI3K signaling in T cell differentiation is underscored by inborn errors of immunity (IEIs) in which altered PI3K signaling drives immune cell dysfunction ^1,2^; activating mutations affecting the catalytic subunit of PI3Kδ cause Activated PI3K delta Syndrome type I (APDS1), an IEI characterized by recurrent respiratory infections, chronic EBV infection, lymphopenia, splenomegaly and lymphadenopathy, including increased Type I responses ^3,4^. Recently, it has emerged that APDS patients show a range of other clinical presentations, including Th2-driven pathologies, such as eosinophilic esophagitis, atopic dermatitis and asthma ^5–7^. However, the molecular basis of T cell dysfunction, particularly in CD4^+^ T helper cells, remains incompletely characterized.

Early activation of naïve CD4^+^ T cells results in the production of IL-2 and activation of STAT5, which are critical for Th1 and Th2 differentiation ^8–11^. In addition to STAT5, IL-2 signaling induces activation of PI3K, mechanistic Target of Rapamycin Complex (mTORC) 1 and mitogen activated protein kinases (MAPKs) ^12–15^, which further promote T cell activation. Together, TCR and cytokine signals drive the induction of lineage defining transcription factors (TFs) that specify CD4 T helper cell fate, including Tbet in Th1 cells and GATA3 in Th2 cells, which in turn negatively cross-regulate one another. In addition, numerous negative regulators must be suppressed to permit T cell activation and differentiation. One such TF that is instrumental in maintaining naïve T cell programs is Foxo1 ^16–18^. Following T cell stimulation, Foxo1 is phosphorylated in a PI3K-dependent fashion, leading to its exclusion from the nucleus and targeting for degradation ^19,20^. Foxo1 inhibition switches off the naïve T cell program, allowing activated cells to take on effector characteristics. In concert with these transcriptional changes, extensive epigenetic remodeling occurs during T cell activation, which reorganizes chromatin accessibility and topology to support and maintain T cell lineage adoption ^21^. One potent determinant of 3D chromatin architecture and gene expression is the transcriptional regulator CTCF. In conjunction with cohesin, CTCF guides the formation of topologically associating domains (TADs) and chromatin looping, which has wide ranging consequences for the regulation of gene expression, allowing chromatin conformations that support dynamic changes in T cell effector states ^22^.

Although PI3K signaling has been extensively studied, the interplay between PI3K signaling and the transcriptional and epigenetic changes that occur during CD4 T cell differentiation remain unclear. Activation of PI3K drives the generation of phosphatidylinositol (3,4,5)-trisphosphate (PIP_3_), leading to recruitment and activation of multiple effectors, including AKT. Activation of AKT has numerous consequences, including phosphorylation and inactivation of TFs such as Foxo1. In many cell-types, PI3K and AKT are also intimately connected to the activation of mTORC1, promoting both Th1 and Th17 differentiation, and mTORC2, further activating AKT, and promoting both Th1 and Th2 cells ^23–25^. Accordingly, activated PI3Kδ enhances cytokine production in multiple CD4 T cell lineages ^5,26^. Conversely, *in vitro* polarization assays utilizing PI3K inhibitors or kinase-dead *Pik3cd^D910A/D910A^* CD4 T cells show impaired Th1, Th2 and Th17 differentiation ^27–29^. How and if PI3K differentially regulates molecular signatures in distinct CD4 T helper lineages remains to be explored.

Examining the consequences of activating mutations affecting *PIK3CD* offers a unique opportunity to better understand the role of PI3Kδ in CD4 T cell biology. Here, we utilized a mouse model expressing the most common variant found in APDS (*Pik3cd^E1020K/+^* mice) to examine Th2 responses using house dust mite (HDM) sensitization, a model of allergic airway inflammation. Unexpectedly, we found that hyperactivated PI3Kδ rewires CD4 T cell differentiation with aberrant expression of IFNγ under Th2 inducing conditions both *in vivo* and *in vitro*. This dysregulation was driven by inappropriate IL-2 and PI3Kδ-induced Foxo1 inactivation, causing a loss of Th2 lineage restriction associated with altered chromatin accessibility and global alterations of CTCF-chromatin interactions. Surprisingly, we linked these phenotypes to elevated expression of the Foxo1-repressed gene *Fasl* in *Pik3cd^E1020K/+^* CD4 T cells, and Fas-mediated potentiation of TCR signaling, which was supported by evidence of interactions between Fas and TCR signaling components, revealed by BioID-based proximity labeling. Collectively, these data uncover non-apoptotic Fas-FasL signaling as a critical amplifier of phenotypes driven by activated PI3Kδ, thereby linking two key pathways of immune dysregulation.

## Results

### Activated PI3Kδ reshapes the HDM induced immune response

To evaluate how activated PI3Kδ affects Th2 differentiation *in vivo*, we examined responses to sensitization with house dust mite (HDM), a model of allergic airway inflammation characterized by Th2 cell and eosinophilic infiltrates and airway hyperreactivity (Fig 1A). Histological staining of lung sections revealed increased immune cell infiltration in lungs of HDM-treated *Pik3cd^E1020K/+^* mice relative to WT counterparts, with significantly elevated inducible bronchus associated lymphoid tissue (iBALT), indicative of an enhanced response (Fig. 1B, Fig S1A). Increased CD4^+^ T cells, the primary mediators of HDM induced responses, were also observed in *Pik3cd^E1020K/+^*lungs (Fig. 1C). However, while HDM-treated WT mice showed a mixed infiltrate of lymphocytes, macrophages and eosinophils, *Pik3cd^E1020K/+^*mice had reduced numbers of eosinophils (Fig 1D) and elevated numbers of neutrophils compared to WT (Fig 1E). Lungs from HDM-treated *Pik3cd^E1020K/+^*mice also exhibited a reduction of total mucosal area compared to both naïve *Pik3cd^E1020K/+^* and WT HDM-treated animals, suggestive of an impaired tissue repair response (Fig S1B). HDM-treated WT animals showed robust airway hyperresponsiveness (AHR), with elevated airway resistance (Rrs) in response to increasing concentrations of methacholine (Fig 1F). In contrast, HDM-treated *Pik3cd^E1020K/+^*lungs completely lacked an AHR signature and instead exhibited a pattern of airway resistance similar to unsensitized naïve animals. Thus, HDM induced a robust but distinct response in mice expressing activated PI3Kδ.

**Figure 1.**
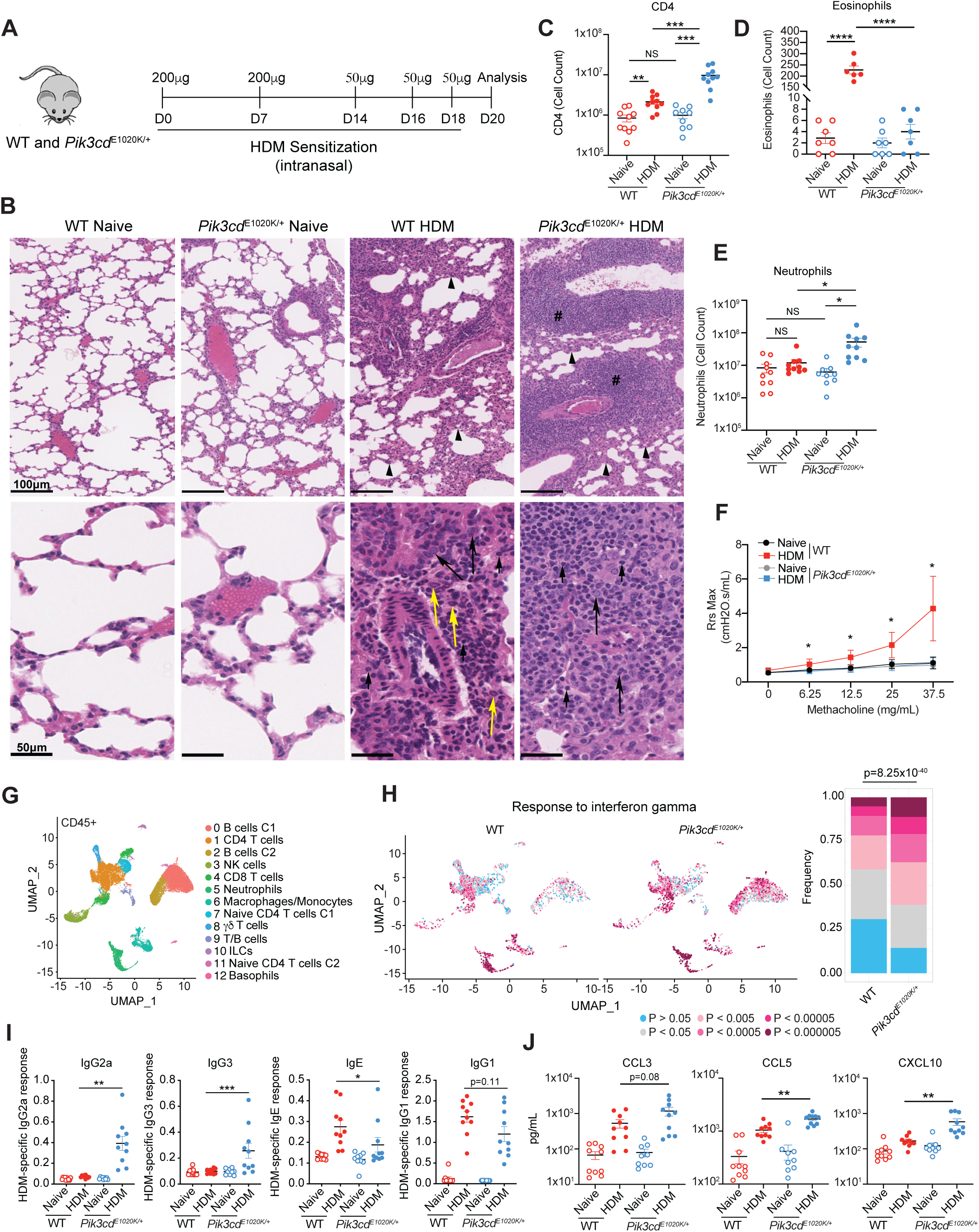
Activated PI3Kδ reshapes the HDM induced immune response. A-J) WT and *Pik3cd^E1020K/+^*animals were sensitized intranasally with HDM extracts (200μg on days 0 and 7; 50μg on days 14, 16 and 18). Mice were euthanized on day 20 and lungs were examined. n=9-10 for each group, pooled from 2 independent experiments. A) Experimental outline. B) Hematoxylin and eosin (H&E) staining of paraffin embedded lung sections from the indicated mice. Top panel, arrowheads indicate thickening of the alveolar septa and pound signs show inducible bronchus associated lymphoid tissue (iBALT). Bottom panel, short arrows show lymphocytes, long arrows show macrophages, yellow arrows show eosinophils. C) Cell counts of CD4 T cells (live CD45^+^TCRβ^+^CD4^+^CD8^-^) measured by flow cytometry. D) Total numbers of eosinophils quantified from H&E stained lung sections. E) Cell counts of neutrophils (liveLineage^-^CD11b^+^Ly6G^+^) from lungs of the indicated animals. F) Airway resistance was quantified from the indicated mice using a methacholine challenge test with a flexivent instrument. Resistance (Rrs) was measured using the indicated concentrations of methacholine as cmH2O.s/mL. Data are representative of 2 independent experiments with n=3-4 for each group per experiment. G) Uniform manifold approximation and projection (UMAP) showing clusters 0-12 of lung CD45^+^ immune cells analyzed by scRNA-seq from naïve WT, naïve *Pik3cd^E1020K/+^*, HDM treated WT and HDM treated *Pik3cd^E1020K/+^* animals. Each cluster was assigned a cell type using singleR and supervised analysis of gene expression specific to each cluster (Supplementary Table 1). Cells from 3 mice were analyzed per genotype and condition. H) Individual lung CD45^+^ immune cells identified by scRNA-seq were analyzed for enrichment of the response to interferon gamma gene set. Left, UMAP visualization of enrichment p-values in lung CD45^+^ immune cells from the indicated mice, with p-values described in the color scale below. Right, frequencies of cells with the indicated magnitudes of enrichment p-values. Statistical comparison of all CD45^+^ cell from WT and *Pik3cd^E1020K/+^* HDM treated groups was performed (Chi-squared test), comparing frequencies of cells with p<0.0005. I) Magnitudes of HDM-specific IgG2a, IgG3, IgE and IgG1 from the indicated mice. J) Chemokine concentrations (pg/mL) of CCL3, CCL5 and CXCL10 measured from lung homogenates from the indicated mice by Luminex. Unless otherwise indicated, statistical comparisons were made using unpaired t-tests. *p<0.05, **p<0.01, ***p<0.001, ****p<0.0001

To better understand how activated PI3Kδ affects responses to HDM, we performed single cell RNA sequencing (scRNAseq) of total CD45^+^ cells from lungs of naïve and HDM-treated mice (Fig 1G, S1C). Unsupervised clustering allowed identification of 13 distinct populations of immune cells, defined by unique gene expression signatures (Supplementary Table 1). For all populations identified, we performed differential gene expression analysis comparing WT and *Pik3cd^E1020K/+^* HDM-treated populations and analyzed genes upregulated in *Pik3cd^E1020K/+^* cells using pathway enrichment analysis. Notably, we saw a prominent enrichment of the response to interferon-gamma gene set as well as a neutrophil chemotaxis signature in multiple populations (Fig S1D). Analysis of individual cells in each cluster revealed increased frequencies of cells with highly significant enrichments (p<0.0005) of response to interferon-gamma signatures in *Pik3cd^E1020K/+^* lungs, including CD4 T cells (Cluster 1), CD8 T cells (Cluster 4), neutrophils (cluster 5), macrophage/monocytes (cluster 6), γδ T cells (cluster 8) and B cells (clusters 0 and 2) (Fig 1H, S1E). Accordingly, serum from *Pik3cd^E1020K/+^* mice showed significantly elevated HDM-specific IgG2a and IgG3 (Fig 1I), which are induced through B cell class switch recombination driven by IFNγ ^30,31^, and are not observed in WT animals following HDM-treatment. Conversely, *Pik3cd^E1020K/+^*serum showed reduced HDM-specific IgE and moderately reduced IgG1 compared to WT (Fig 1I). Lung homogenates from HDM-treated *Pik3cd^E1020K/+^* animals also exhibited elevated concentrations of a number of IFNγ-induced chemokines, including CCL3, CCL5 and CXCL10 (Fig 1J). Therefore, *Pik3cd^E1020K/+^* mice showed a global change in polarized responses to HDM, reshaping immunity towards a type I inflammatory response.

### Profound Th1 responses in *Pik3cd^E1020K/+^* lungs following house dust mite sensitization, at the expense of Th2 immunity

To further understand these phenotypes, we focused on CD4 T cells, which are the primary mediators of responses to HDM. Differential gene expression analysis comparing HDM-treated WT and *Pik3cd^E1020K/+^* CD4 T cells in pseudobulk (cluster 1, Figure 1G) revealed reduced expression of genes characteristic of pathogenic Th2 cells in *Pik3cd^E1020K/+^*CD4 cells, including *Gata3*, *Il1rl1,* encoding the IL-33 receptor ST2, and *Areg*, encoding Amphiregulin, a protein associated with tissue repair in asthma (Fig 2A); UMAP projections of total CD45^+^ cells confirmed a near absence of expression of these and other key Th2 lineage defining genes (Fig S2A). In contrast, we observed increased expression of genes associated with a cytolytic Th1 signature in *Pik3cd^E1020K/+^* CD4 cells compared to those from WT mice, including *Ifng,* as well as *Gzmb* and *Nkg7* (Fig 2A).

**Figure 2.**
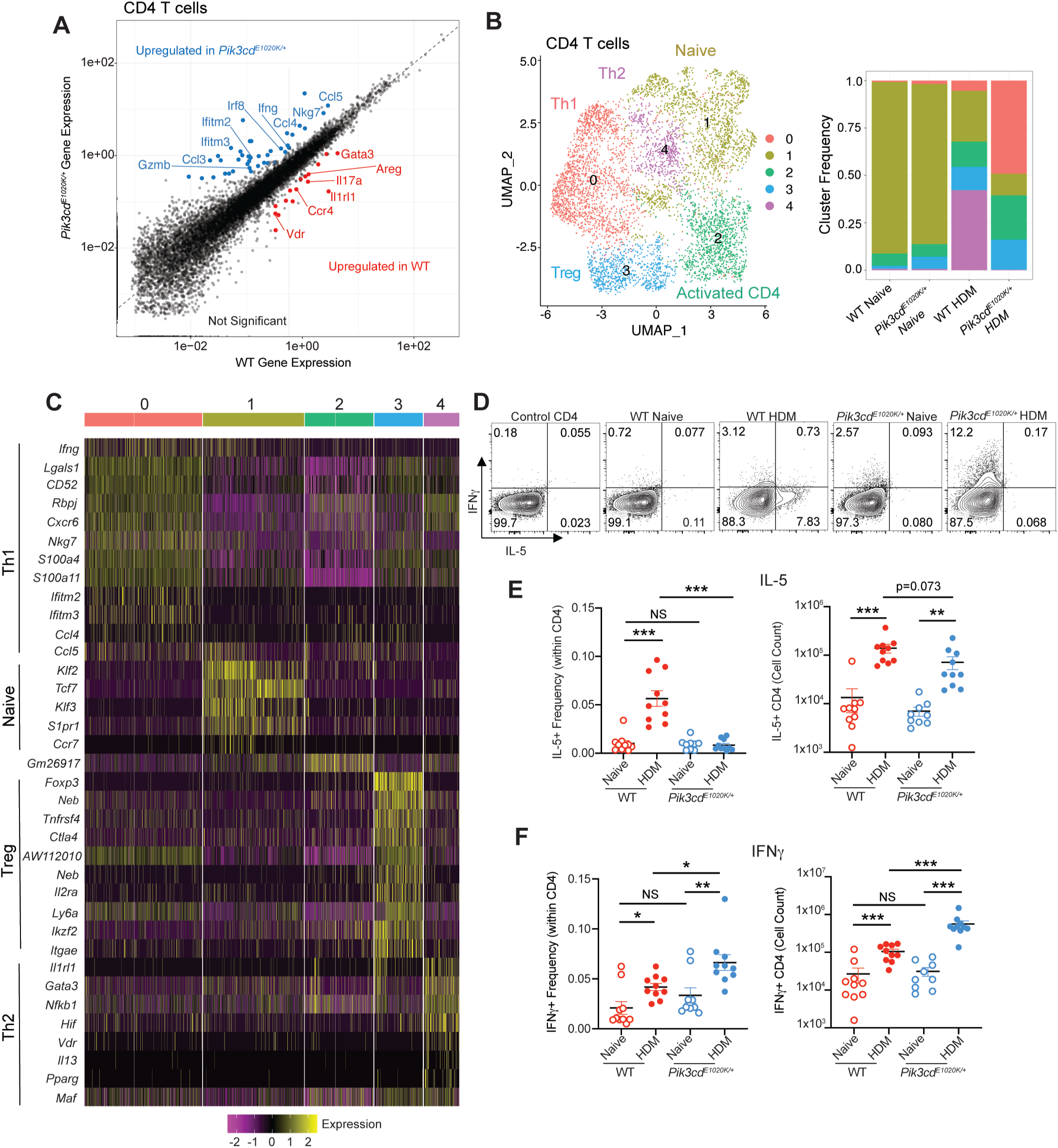
Aberrant Th1 responses at the expense of Th2 immunity in *Pik3cd^E1020K/+^* lungs following house dust mite sensitization. A) Scatter plot comparing gene expression in WT and *Pik3cd^E1020K/+^* HDM treated CD4 T cells (Cluster 1 of total CD45^+^ in Fig 1G). Differentially expressed genes are highlighted; genes upregulated in WT CD4 T cells are highlighted in red, genes upregulated in *Pik3cd^E1020K/+^* CD4 T cells are highlighted in blue. B) CD4^+^ T cells (cluster 1 of total CD45^+^) were identified from scRNA-seq of total CD45^+^ lung immune cells from naïve WT, naïve *Pik3cd^E1020K/+^*, HDM treated WT and HDM treated *Pik3cd^E1020K/+^* animals and re-clustered to identify 5 unique clusters (0-4) of CD4^+^ T cells. Left, UMAP showing distribution of clusters 0-4; identities of each cluster were assigned based on cluster specific gene expression and are indicated. Right, proportion of cells from each cluster from the indicated mice. C) Seurat heatmap showing expression of cluster defining genes for the indicated populations. D) Representative flow cytometry plots of lung CD4^+^ T cells (liveCD45^+^TCRβ^+^CD4^+^CD8^-^) measuring intracellular IFNγ and IL-5 expression in cells from the indicated mice. n=9-10 for each group, pooled from 2 independent experiments. E) Frequencies (left) and cell counts (right) of IL-5^+^ lung CD4 T cells from the indicated groups. F) Frequencies (left) and cell counts (right) of IFNγ^+^ lung CD4 T cells from the indicated groups. Statistical comparisons were made using unpaired t-tests. *p<0.05, **p<0.01, ***p<0.001.

We next reclustered CD4 T cells (cluster 1, Fig 1G) alone, identifying 5 distinct sub-populations defined by unique gene expression patterns (Fig 2B, C). Cluster 1 showed high expression of signature naïve CD4 T cell genes such as *Ccr7*, *S1p1r* and *Klf2*, and was prominent in both naïve WT and *Pik3cd^E1020K/+^* animals but reduced after HDM-treatment (Fig 2B, C). Cluster 4, which was defined by expression of *Gata3*, *Il1rl1* and *Il13* as a Th2 population (Fig 2C) was increased in WT CD4 cells post-HDM. However, this cluster was virtually absent among *Pik3cd^E1020K/+^* CD4 T cells (Fig 2B). In contrast, cluster 0, which was defined by expression of *Ifng* and numerous Th1 signature genes, dominated amongst *Pik3cd^E1020K^* CD4 T cells post-HDM, despite making up only a minor fraction of WT CD4 cells.

Flow cytometry confirmed that HDM induced a population of CD4 cells expressing GATA3 in WT lungs; however, these were significantly reduced, both in number and frequency, in lungs of *Pik3cd^E1020K/+^* mice (Fig S2B). Similarly, Th2 cytokines, IL-4, IL-5 and IL-13 were produced in WT CD4 cells in response to HDM, but were markedly diminished in frequency in CD4 T cells from *Pik3cd^E1020K/+^* mice (Fig 2D, E and S2C). In contrast, HDM-treated *Pik3cd^E1020K/+^* CD4 showed elevated numbers and frequencies of IFNγ-producing cells relative to WT (Fig 2D, F). Cytokine measurements from total lung homogenates confirmed significantly impaired induction of type 2 cytokines IL-4 and IL-5 but elevated concentrations of TNFα and GM-CSF in *Pik3cd^E1020K/+^* mice (Fig S2D, E). Thus, *Pik3cd^E1020K/+^*CD4^+^ T cells showed a profound switch towards Th1 differentiation at the expense of Th2 lineage adoption.

To evaluate whether these phenotypes were CD4 T cell intrinsic, naïve CD4 T cells from WT or *Pik3cd^E1020K/+^* mice were adoptively transferred into TCRα-deficient recipients, which were then sensitized with HDM (Fig S3A). Similar to intact animals, mice receiving *Pik3cd^E1020K/+^* CD4 T cells showed elevated numbers of lung CD4 T cells following HDM-treatment, along with impaired induction of GATA3 expression, diminished frequencies of CD4 cells producing Type 2 cytokines (IL-4, 5, 13) and elevated frequencies of IFNγ-producing CD4 cells (Fig S3B-D). The defective Type 2 responses to HDM in *Pik3cd^E1020K/+^* mice were, therefore, CD4 T cell-intrinsic.

### Loss of Th2 lineage restriction in *Pik3cd^E1020K/+^* CD4 T cells

To probe CD4^+^ T cell differentiation in a defined system, we examined cytokine production from isolated naïve CD4 cells that were differentiated *in vitro* in the presence of WT APCs. In contrast to *in vivo* responses to HDM, *in vitro* Th2 differentiation of *Pik3cd^E1020K/+^* CD4 cells gave rise to increased frequencies of Th2 cytokine producing cells, including IL-4^+^ and IL-13^+^ cells, along with intact expression of GATA3 (Fig S3E, F). However, *Pik3cd^E1020K/+^* Th2 polarized cells also exhibited significantly elevated frequencies of IFNγ-expressing cells relative to WT Th2 cells, which showed minimal IFNγ production (Fig 3A). Moreover, *Pik3cd^E1020K/+^* Th2 cells expressed elevated Tbet, similar to that seen in WT Th1 cells (Fig 3B), despite normal expression of GATA3 (Fig S3F). Increased IFNγ expression could be detected as early as 24h into differentiation, with frequencies of IFNγ^+^ cells steadily increasing over time in *Pik3cd^E1020K/+^* Th2 cells (Fig S3G) and could be prevented by anti-IFNγ blocking antibodies (Fig S3H). Thus, cells expressing activated PI3Kδ showed a loss of Th2 lineage restriction.

**Figure 3.**
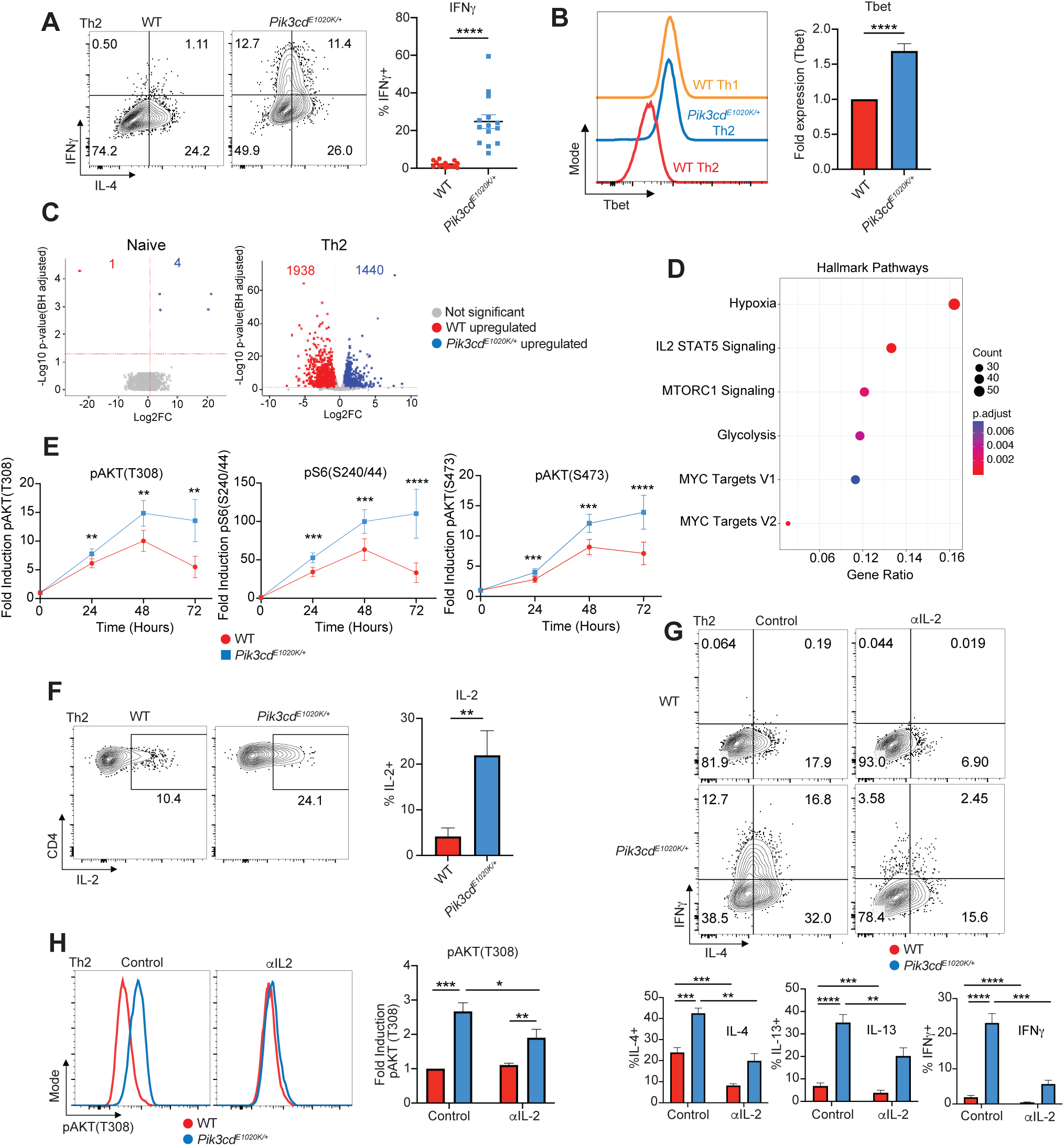
Hyperactivated PI3Kδ disrupts Th2 lineage restriction through enhanced IL-2 signaling. A-D) Naïve CD4 T cells were activated with aCD3 + aCD28 in the presence of WT APCs under Th2 polarizing conditions (IL-4 + aIL-12) for 72 hours. n=14 for each group, 14 independent experiments. A) Left, Representative flow cytometry plots showing IFNγ and IL-4 staining in Th2 polarized live CD4^+^ cells. Right, Percentages of IFNγ^+^ cells from the indicated mice. B) Left, representative flow cytometry histograms showing Tbet expression in Th2 and WT control Th1 polarized live CD4^+^ cells from the indicated mice. Right, Fold Tbet expression (MFI normalized to WT) in Th2 polarized live CD4^+^ cells from the indicated mice. C) Volcano plots showing differentially expressed genes (DEGs) in red (WT upregulated) and blue (*Pik3cd^E1020K/+^*upregulated); DEGs defined using fold change>1.5, p<0.05. Bulk RNAseq was performed on WT and *Pik3cd^E1020K/+^* naïve CD4 T cells and *in vitro* polarized Th2 cells. D) Enrichment of hallmark pathways among DEGs comparing WT and *Pik3cd^E1020K/+^* Th2 cells. E) Fold induction of pAKT(T308), pS6(S240/44) and pAKT(S473) in WT and *Pik3cd^E1020K/+^* live CD4^+^ cells over a time course of Th2 differentiation (0, 24, 48 and 72 hours), measured by flow cytometry. Fold induction was calculated using MFIs of the indicated readouts normalized to the 0 timepoint of the corresponding genotype. n=5-9 for each group, 5-9 independent experiments. F) Left, Representative flow cytometry plots showing IL-2 and CD4 expression in Th2 polarized cells from the indicated mice. Right, percentages of IL-2^+^ Th2 cells from the indicated mice. n=9 for each group, 9 independent experiments. G-H) Naïve CD4 T cells from the indicated mice were Th2 polarized in the presence or absence of αIL-2 blocking antibody. G) Top, representative flow cytometry plots showing IFNγ and IL-4 staining in Th2 polarized live CD4^+^ cells. Bottom, Percentages of IL-4^+^ (left), IL-13^+^ (middle) and IFNγ^+^ (right) cells from the indicated mice, in the presence or absence of αIL-2. n=11 for each group, 11 independent experiments. H) Left, Representative flow cytometry plots showing pAKT(T308) in Th2 polarized live CD4^+^ cells from the indicated groups. Right, fold induction of pAKT(T308) in Th2 polarized live CD4^+^ cells from the indicated groups. Fold induction was calculated by normalizing pAKT(T308) MFIs to WT control cells. n=5 for each group, 5 independent experiments. Statistical comparisons were made using ratio paired t-tests. *p<0.05, **p<0.01, ***p<0.001, ****p<0.0001.

### Elevated IL-2-STAT5 signaling in *Pik3cd^E1020K/+^* cells drives aberrant Th2 differentiation and heightened PI3K-AKT-mTOR signaling

We next performed bulk RNAseq analysis of *in vitro* polarized Th2 cells from WT and *Pik3cd^E1020K/+^* mice. Transcriptomic analyses revealed remarkably few differentially expressed genes (DEGs) between WT and *Pik3cd^E1020K/+^*naïve CD4 T cells (Fig 3C). However, comparison of day 3 differentiated WT and *Pik3cd^E1020K/+^* Th2 cells, showed several thousand DEGs (Fig 3C), highlighting the profound consequences of PI3Kδ hyperactivation on gene expression following Th2 polarization.

Pathway enrichment analysis using Th2 specific DEGs revealed an enrichment of the mTORC1 and MYC signaling pathways (Fig 3D). This was confirmed by time course analysis of PI3K-Akt and mTOR signaling (Fig 3E); *Pik3cd^E1020K/+^*Th2 cells showed significantly elevated induction of pAKT(T308) following stimulation, as well as increased activation of both mTORC1, as evidenced by increased phosphorylation of ribosomal protein S6 (pS6(S240/44)), and mTORC2, as evidenced by increased phosphorylation of AKT on Ser473, compared to WT (Fig 3E). Additionally, enrichment of IL-2-STAT5 signaling pathways was observed among Th2 DEGs (Fig 3D); this corresponded to increased frequencies of IL-2^+^ *Pik3cd^E1020K/+^* differentiated Th2 cells, with *Pik3cd^E1020K/+^* Th2 cells showing sustained IL-2 production compared to WT counterparts (Fig 3F, S3I). Accordingly, *Pik3cd^E1020K/+^*cells showed elevated and sustained pSTAT5(Y694) relative to WT following differentiation (Fig S3J).

To determine whether *Pik3cd^E1020K/+^* cells also respond more potently to IL-2 stimulation, cells were differentiated under Th2 conditions, rested in cytokine-free media and then stimulated with IL-2 and evaluated for STAT5 and S6 phosphorylation. WT and *Pik3cd^E1020K/+^*cells had similar surface CD25, ruling out altered IL-2Rα (CD25) expression as a possible variable (Fig S3K). Although pSTAT5 was similar between WT and *Pik3cd^E1020K/+^*cells (Fig S3L), *Pik3cd^E1020K/+^* Th2 cells showed increased pS6(S240/44) over the course of IL-2 stimulation (Fig S3M), suggesting that *Pik3cd^E1020K/+^* Th2 cells both produce more IL-2 and respond more strongly to IL-2 stimulation, with increased mTORC1 activity.

To determine whether IL-2-mediated pathways contributed to altered CD4 T cell differentiation, we performed *in vitro* differentiation experiments in the presence or absence of an IL-2 blocking antibody (αIL-2) (Fig 3G, H). Both WT and *Pik3cd^E1020K/+^* Th2 polarized cells showed significant reductions in frequencies of IL-4^+^ and IL-13^+^ cells following aIL-2 treatment, consistent with roles for IL-2 in promoting Th2 differentiation (Fig 3G). However, inappropriate IFNγ production by *Pik3cd^E1020K/+^* Th2 polarized cells was markedly reduced by blockade of IL-2, with aIL-2 treated *Pik3cd^E1020K/+^*Th2 showing significantly depressed frequencies of IFNγ^+^ cells compared to control (Fig 3G). IL-2 blockade also diminished the induction of pAKT(T308) in *Pik3cd^E1020K/+^*Th2 cells (Fig 3H), suggesting that IL-2 exacerbates PI3Kδ signaling in these cells.

### IL-2 represses Foxo1 activity in *Pik3cd^E1020K/+^* CD4^+^ T cells

To elucidate potential transcription factors regulated by activated PI3Kδ during Th2 differentiation, we performed gene set enrichment analysis (GSEA) comparing WT and *Pik3cd^E1020K/+^* cells for transcription factor target (TFT) gene expression (Fig S4A). We observed significant enrichment of many more TFT gene sets in WT cells compared to *Pik3cd^E1020K/+^*including AP4, MYOD, MEIS1 as well as genes regulated by Foxo1, a key transcription factor that is inactivated by phosphorylation via Akt (Fig S4A). Notably, analysis of DEGs upregulated in *Pik3cd^E1020K/+^*CD4 T cells following HDM-treatment *in vivo* also revealed significant enrichment of a Foxo1-knockout signature (Fig 4A). TCR-mediated activation of PI3Kδ leads to phosphorylation and inactivation of Foxo1; pFoxo1 was observed in response to TCR stimulation in both WT and *Pik3cd^E1020K/+^*CD4 cells following T cell activation (Fig 4B). However, while Foxo1 phosphorylation subsided in WT Th2 cells by 72h of stimulation, pFoxo1(S256) remained high in *Pik3cd^E1020K/+^* cells. Accordingly, we observed decreased total Foxo1 protein in *Pik3cd^E1020K/+^* Th2 cells over the differentiation period (Fig S4B).

**Figure 4.**
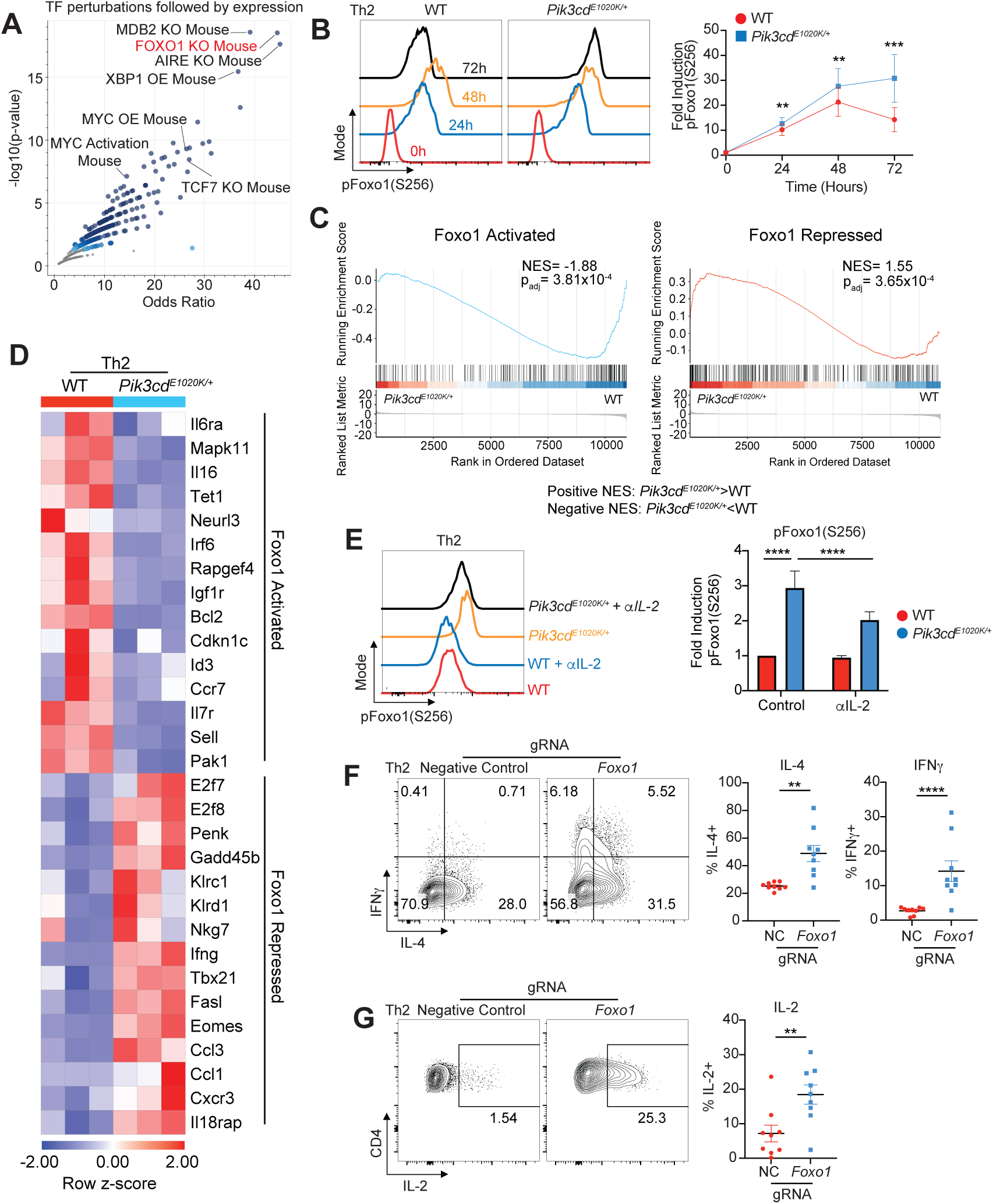
IL-2 driven inactivation of Foxo1 in *Pik3cd^E1020K/+^* CD4^+^ T cells impairs Th2 lineage restriction. A) Pathway enrichment of TF Perturbations Followed by Expression gene sets was performed using Enrichr ^73^, with dots representing significantly enriched gene sets colored in blue; genes upregulated HDM treated *Pik3cd^E1020K/+^* CD4 T cells relative to WT counterparts (Fig 2A) were used as input for pathway enrichment. B) Time course analysis (0, 24, 48, 72 hours) of pFoxo1(S256) was performed using Th2 polarized cells from the indicated mice. Left, Representative flow cytometry plots showing pFoxo1(S256) over time in Th2 polarized live CD4^+^ cells from the indicated groups. Right, fold induction of pFoxo1(S256) over time in Th2 polarized live CD4^+^ cells from the indicated groups. Fold induction was calculated by normalizing pFoxo1(S256) MFIs to the 0 time point of the corresponding genotype. n=8 for each group, 8 independent experiments. C) Gene set enrichment analysis (GSEA) comparing WT and *Pik3cd^E1020K/+^* transcriptomes for the expression of Foxo1 activated (left) and repressed (right) gene sets. D) Gene expression heatmap (row z-score) showing normalized RPKM values of leading edges from Foxo1 activated and repressed GSEA described in figure 4C. E) Naïve CD4^+^ T cells from the indicated mice were Th2 polarized in the presence or absence of αIL-2 blocking antibody. Left, Representative flow cytometry plots showing pFoxo1(S256) in Th2 polarized live CD4^+^ cells from the indicated groups. Right, fold induction of pFoxo1(S256) in Th2 polarized live CD4^+^ cells from the indicated groups. Fold induction was calculated by normalizing pFoxo1(S256) MFIs to WT control cells. n=10 for each group, 10 independent experiments. F-G) Naïve CD4 T cells were nucleofected with Cas9-gRNA complexes containing negative control or *Foxo1* targeting gRNAs and differentiated under Th2 conditions. n=13 for each group, 13 independent experiments. F) Left, Representative flow cytometry plots showing IFNγ and IL-4 expression in live CD4^+^ T cells from the indicated groups. Right, percentages of IFNγ^+^ and IL-4^+^ cells in Th2 polarized cells from the indicated mice. G) Left, Representative flow cytometry plots showing IL-2 and CD4 expression in live CD4^+^ T cells from the indicated groups. Right, percentages of IL-2^+^ cells in Th2 polarized cells from the indicated groups. Statistical comparisons were made using ratio paired t-tests. **p<0.01, ***p<0.001, ****p<0.0001.

Using publicly available transcriptomic data, we generated lists of known Foxo1 activated and repressed genes ^32^ and compared transcriptomes of WT versus *Pik3cd^E1020K/+^* cells by GSEA (Fig 4C, Supplementary Table 2). While WT cells exhibited significantly enriched expression of Foxo1 activated genes, *Pik3cd^E1020K/+^* cells showed significant enrichment of Foxo1 repressed gene signatures relative to WT (Fig 4C). Focusing on leading-edge genes identified by GSEA (Fig 4D), we found multiple relevant Foxo1 activated genes were enriched in WT cells including *Sell*, *Ccr7*, *Il7r* and *Bcl2*. Conversely, Foxo1-repressed genes that were enriched in *Pik3cd^E1020K/+^* cells had a notable type I immune response signature, including *Ifng*, *Tbx21*, *Eomes* and *Fasl*.

To evaluate whether IL-2 signaling influenced Foxo1 activity, we measured pFoxo1(S256) following differentiation in the presence or absence of an IL-2 blocking antibody (Fig 4E). Although αIL-2 treatment had little effect on pFoxo1(S256) induction in WT cells, *Pik3cd^E1020K/+^* cells showed reduced pFoxo1(S256) in αIL-2 conditions compared to control. In contrast, blocking IFNγ did not significantly alter phosphorylation of Foxo1 (Fig S4D), although it decreased aberrantly high expression of Tbet and IFNγ in *Pik3cd^E1020K/+^* Th2 cells (Fig S3H).

We next examined whether IL-2 signaling regulates Foxo1 transcriptional activity. GSEA comparison of transcriptomes of Th2 cells differentiated in the presence or absence of αIL-2 (Fig S4C) showed significant enrichment of Foxo1-activated gene signatures in both αIL-2 treated WT and *Pik3cd^E1020K/+^*Th2 cells, compared to controls. Conversely, both control WT and *Pik3cd^E1020K/+^*Th2 cells showed significant enrichment of Foxo1-repressed genes relative to αIL-2 treated cells under Th2 conditions, suggesting that IL-2 is a key inhibitor of Foxo1 transcriptional activity in CD4 T cells.

### Foxo1 is critical for Th2 lineage restriction

To address whether Foxo1 directly contributes to altered CD4 T cell differentiation, we targeted *Foxo1* using Cas9 ribonucleoprotein (RNP) complexes in naïve CD4 T cells prior to *in vitro* Th2 polarization (Fig S4E). *Foxo1*-targeted Th2 polarized cells showed elevated frequencies of IL-4^+^ cells, relative to negative controls (Fig 4F). However, Foxo1-deficient Th2 cells also showed aberrant production of IFNγ, recapitulating the phenotype of *Pik3cd^E1020K/+^*polarized cells. Conversely, retroviral-mediated expression of Foxo1-WT or Foxo1-AAA, a Foxo1 mutant resistant to AKT-mediated phosphorylation, decreased Th2 cytokine production, especially in WT cells (Fig S4F). Notably, aberrant IFNγ production by *Pik3cd^E1020K/+^* Th2 polarized cells was markedly reduced by either Foxo1-WT or Foxo1-AAA overexpression, with frequencies of IFNγ^+^ cells significantly reduced compared to control cells. Thus, Foxo1 activity appears to be critical to prevent inappropriate IFNγ expression in Th2 cells. Foxo1-deficient Th2 cells also showed increased frequencies of IL-2^+^ cells compared to controls (Fig 4G). Therefore, Foxo1 both suppresses IL-2 expression and is suppressed by IL-2, highlighting a signaling amplification loop whereby increased IL-2 production promotes further Foxo1 phosphorylation and inactivation.

### *Pik3cd^E1020K^* reshapes the epigenetic landscape of CD4^+^ T cells

Activation and differentiation of CD4 cells are associated with dynamic global changes to chromatin architecture that are required for CD4 T cell differentiation. To probe the downstream consequences of this amplified PI3Kδ-Foxo1-IL-2 signaling circuit, we performed ATACseq analysis of *in vitro* differentiated WT and *Pik3cd^E1020K/+^* Th2 cells. We observed differential accessibly of ∼13% of the epigenome (Fig 5A, S5A). Among these differential regions, we sought to identify transcription factor motifs that influence CD4 T cell differentiation. Major transcription factor motif families that were distinct among *Pik3cd^E1020K/+^* Th2 specific peaks included those binding NFAT:AP1 complexes, interferon regulatory factors (IRFs) and runt related transcription factors (Runx), all of which are associated with T cell activation and type I responses (Fig 5B). *Pik3cd^E1020K/+^* Th2 specific peaks further showed increased accessibility of T-box family motifs, including Tbet-binding motifs, reflecting their altered cytokine expression. In contrast, WT Th2 specific peaks showed enrichment of numerous motifs associated with Th2- promoting factors, including STAT6 and GATA3 (Fig 5B). However, in addition to these lineage- and activation-defining transcription factors, CTCF motifs were also differentially accessible, with a strong enrichment in WT Th2 specific peaks. CTCF functions as a global regulator of chromatin organization, suggesting broad alterations associated with activated PI3Kδ.

**Figure 5.**
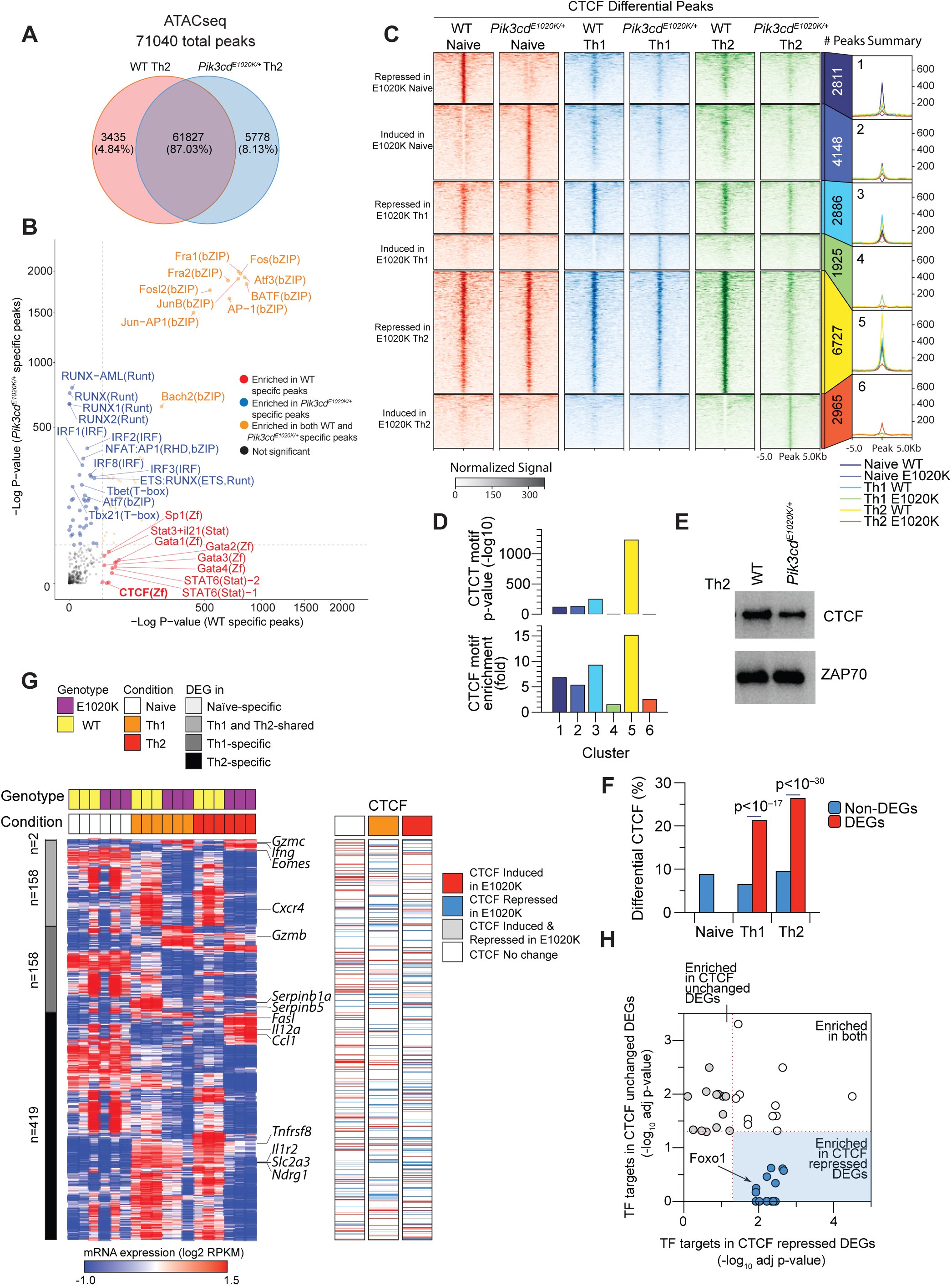
*Pik3cd^E1020K^* reshapes the epigenetic landscape of CD4^+^ T cells. A) Naïve CD4 T cells from WT and *Pik3cd^E1020K/+^* mice were polarized under Th2 conditions and evaluated by ATACseq. A total of 71040 peaks were detected and WT specific, *Pik3cd^E1020K/+^* specific and common peaks are described in a Venn diagram. B) WT and *Pik3cd^E1020K/+^* specific peaks were examined by motif enrichment analysis and enrichment p-values were plotted for both groups. Motifs specifically enriched in WT peaks are indicated in red, motifs specifically enriched in *Pik3cd^E1020K/+^* are indicated in blue and motifs enriched in both groups are indicated in orange. C) Peak heatmap of CTCF CUT&Tag peaks from WT and *Pik3cd^E1020K/+^* naïve, Th1 and Th2 cells; peaks are organized into 6 clusters (1-6) specific to each population, as described. D) CTCF motif enrichment p-value (top) and fold enrichment (bottom) in clusters 1-6. E) Western blot evaluating CTCF and Zap70 in lysates from WT and *Pik3cd^E1020K/+^* Th2 polarized cells. Data are representative of 3 independent experiments. F) Using bulk RNAseq data (Figure 3C), DEGs (WT vs *Pik3cd^E1020K/+^*) and non-DEGs were compared in the indicated populations for percentages of genes showing differential CTCF peaks (WT vs *Pik3cd^E1020K/+^*). G) Left, gene expression heatmap showing RPKM values of DEGs (WT vs *Pik3cd^E1020K/+^*; fold change>4, p<0.05) in naïve, Th1 and Th2 polarized cells. Right, CTCF heatmap aligned with DEG heatmap showing CTCF peaks induced, repressed, both induced and repressed, or with no change in *Pik3cd^E1020K/+^* cells near DEGs. H) Th2 DEGs from figure 5G were organized into two categories; DEGs showing no change in CTCF (WT vs *Pik3cd^E1020K/+^*) and DEGs showing repressed CTCF peaks in *Pik3cd^E1020K/+^* relative to WT. Pathway enrichment analysis (Enrichr ^73^) of transcription factor targets (ChEA ^89^) was performed using these two categories of DEGs and adjusted p-values from the top 25 enriched TF signatures in each category were plotted against each other in the shown scatter plot. Foxo1 targets were specifically enriched among DEGs with repressed CTCF in *Pik3cd^E1020K/+^* cells.

We next employed CUT&Tag to specifically examine genome-wide CTCF binding in WT and *Pik3cd^E1020K/+^* naïve CD4, Th2, and Th1 cells for comparison. Differential CTCF peaks, defined as at least 4-fold difference in peak intensity, were grouped into six clusters, with each cluster representing differential CTCF peaks between WT and *Pik3cd^E1020K/+^*CD4 cells in naïve, Th1 and Th2 polarized states (Fig 5C). We identified approximately 7000, 5000 and 10000 differential CTCF peaks in the naïve, Th1 and Th2 states, respectively, indicating significant genome-wide differences in CTCF binding in *Pik3cd^E1020K/+^* CD4 cells. To further explore these observations, we quantified the enrichment of CTCF motifs within the peaks in each cluster. In naïve cell clusters (clusters 1-2), CTCF motif enrichment showed a roughly equivalent magnitude between WT and *Pik3cd^E1020K/+^* cells (Fig 5D), likely reflecting CTCF repositioning in this state. WT Th1 (cluster 3) and Th2 (cluster 5) specific peaks displayed strong CTCF motif enrichment (Fig 5D). In contrast, the *Pik3cd^E1020K/+^* Th1 and Th2-specific peaks (clusters 4 and 6, respectively) exhibited surprisingly low CTCF motif enrichment (Fig 5D). These findings suggested a loss of CTCF motif recognition in *Pik3cd^E1020K/+^* CD4 T cells, prompting us to assess CTCF protein expression itself (Fig 5E). *Pik3cd^E1020K/+^*Th2 cells demonstrated reduced CTCF protein levels compared to WT counterparts (Fig 5E, S5B), suggesting that activated PI3Kδ negatively regulates CTCF protein expression, leading to widespread impairment of CTCF-DNA interactions in *Pik3cd^E1020K/+^* Th2 cells.

To further explore the relationship between CTCF and gene expression, we annotated differential CTCF peaks (Fig 5C) to genes and compared those associated with DEGs to non-DEGs of WT versus *Pik3cd^E1020K/+^*Th2 cells (Fig 5F). We observed that DEGs showed significantly higher percentages of genes associated with differential CTCF peaks relative to non-DEGs, indicating changes in CTCF profiles at specific loci related to differential gene expression. We next considered CTCF peaks near DEGs and determined the pattern of CTCF binding in the vicinity of these genes (Fig 5G). Consistent with previously seen diminished CTCF motif enrichment (Fig 5D), the bulk of DEGs with differential CTCF peaks demonstrated a loss of CTCF binding in *Pik3cd^E1020K/+^* Th2 cells (Fig. 5G). To determine whether particular TF signatures were associated with repressed CTCF in *Pik3cd^E1020K/+^*cells, we performed pathway enrichment analysis comparing transcription factor targets enriched in DEGs associated with repressed CTCF peaks in *Pik3cd^E1020K/+^*Th2 cells to DEGs showing no change in CTCF (Fig 5H). Notably, DEGs associated with repressed CTCF in *Pik3cd^E1020K/+^* Th2 cells showed significant enrichment of targets for several TFs (Supplementary Table 3), including Foxo1, which were not observed in DEGs with no change in CTCF; analysis of genomic regions surrounding multiple Foxo1 activated and repressed loci revealed altered patterns of CTCF binding in *Pik3cd^E1020K/+^* Th2 cells (Fig S5C). Thus, activated PI3Kδ was associated with diminished CTCF protein and repressed CTCF profiles enriched near multiple DEGs, including those regulated by Foxo1, in *Pik3cd^E1020K/+^*Th2 cells.

### Fas-FasL signaling drives PI3K**δ** hyperactivation in Th2 cells

To provide further insight into how Foxo1 affects Th2 lineage restriction, we explored specific Foxo1 regulated genes. Among these, we noted that *Fasl* was significantly elevated in *Pik3cd^E1020K/+^* CD4 T cells (Fig 4D); increased expression of FasL was confirmed by surface staining and flow cytometry of *Pik3cd^E1020K/+^* Th2 cells (Fig 6A). *Foxo1* gRNA targeted Th2 cells also showed elevated surface FasL expression compared to negative control gRNA treated cells (Fig 6B) whereas αIL-2 treated *Pik3cd^E1020K/+^* Th2 cells showed diminished FasL compared to control counterparts (Fig 6C). Expression of FasL in *Pik3cd^E1020K/+^* Th2 cells therefore paralleled the patterns observed for IFNγ.

**Figure 6.**
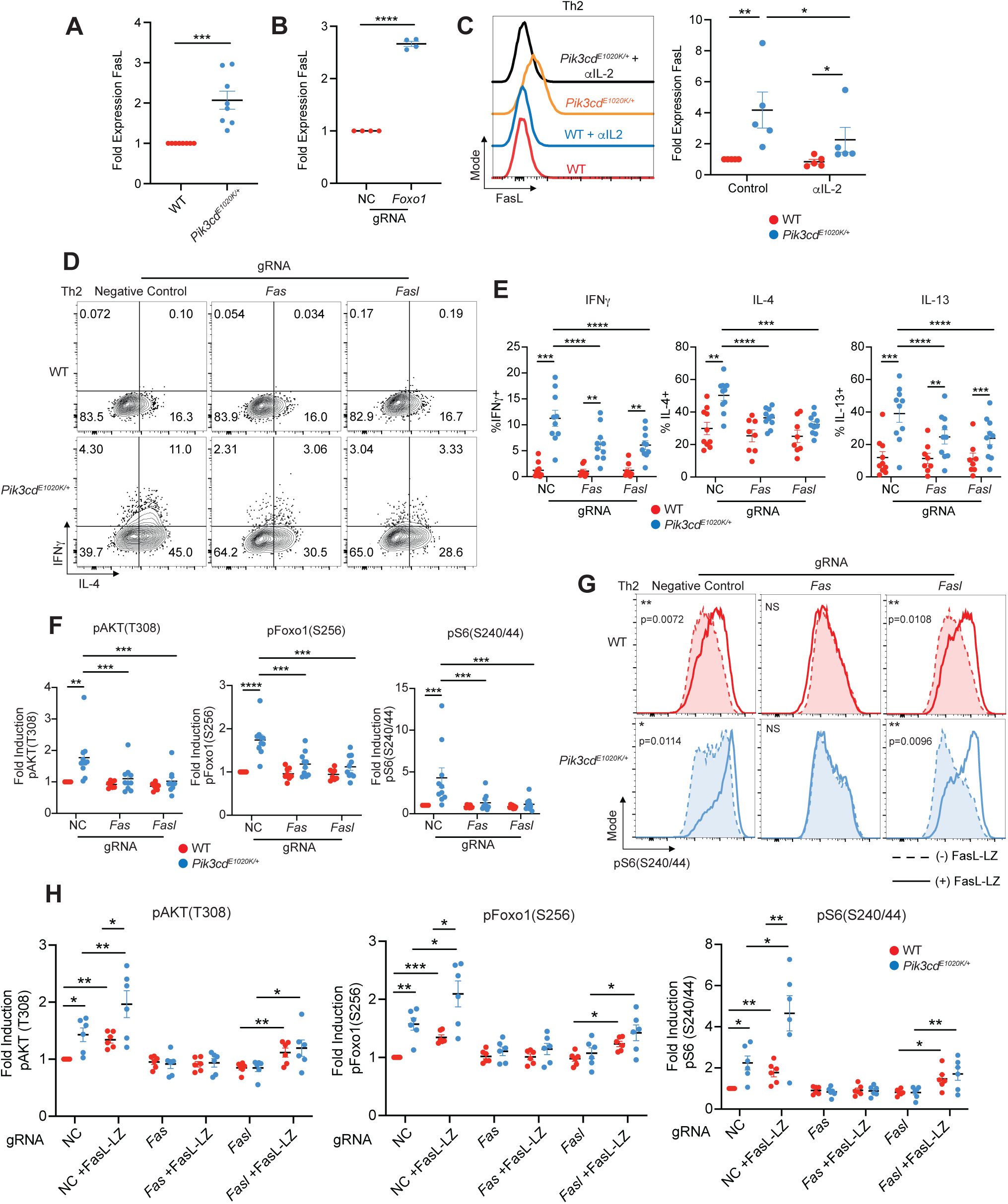
Fas-FasL signaling potentiates T cell activation, exacerbating CD4^+^ T cell dysregulation in APDS. A) Fold FasL expression (MFI normalized to WT) on the surface of Th2 polarized live CD4^+^ T cells, measured by flow cytometry from the indicated mice. n=8 for each group, 8 independent experiments B) Fold FasL expression (MFI normalized to negative control) on the surface of negative control (NC) and *Foxo1* gRNA targeted naïve CD4 T cells cultured under Th2 polarizing conditions. n=4 for each group, 4 independent experiments. C) Naïve CD4^+^ T cells from the indicated mice were Th2 polarized in the presence or absence of αIL-2 blocking antibody and surface FasL expression was measured by flow cytometry. Left, representative flow cytometry histograms of surface FasL. Right, fold FasL expression (MFI normalized to WT control) on Th2 polarized cells from the indicated groups. n=5 for each group, 5 independent experiments. D-F) Naïve CD4 T cells from WT and *Pik3cd^E1020K/+^*mice were nucleofected with Cas9-gRNA complexes containing negative control, *Fas* or *Fasl* targeting gRNAs and underwent Th2 polarization. n=8-10 for each group, 8-10 independent experiments. D) Representative flow cytometry plots showing IFNγ and IL-4 expression in live CD4^+^ T cells from the indicated groups. E) Percentages of IFNγ^+^ (left), IL-4^+^ (middle) and IL-13^+^ (right) Th2 polarized live CD4^+^ T cells from the indicated groups. F) Fold induction of pAKT(T308), pFoxo1(S256) and pS6(S240/44) in Th2 polarized liveCD4 T cells from the indicated groups, measured by flow cytometry. For all readouts, fold induction was calculated by normalizing MFIs to negative control WT cells. G-H) Negative control, *Fas* and *Fasl* gRNA treated naïve CD4 T cells underwent Th2 polarization in the presence or absence of recombinant multimeric FasL (FasL-LZ) and phosphorylation of AKT and S6 were analyzed by flow cytometry. n=6 for each group, 6 independent experiments. G) Representative flow cytometry histograms showing pS6(S240/44) staining in Th2 polarized live CD4^+^ T cells from the indicated groups. Dashed lines represent cells cultured without FasL-LZ, solid lines show cells cultured with FasL-LZ. H) Fold induction of pAKT(T308), pFoxo1(S256) and pS6(S240/44) in Th2 polarized (-/+ FasL-LZ) live CD4^+^ T cells from the indicated groups, measured by flow cytometry. For all readouts, fold induction was calculated by normalizing MFIs to negative control WT cells. Statistical comparisons were made using ratio paired t-tests. *p<0.05, **p<0.01, ***p<0.001, ****p<0.0001.

Although Fas-FasL signaling is best known for its role in modulating apoptosis, non-apoptotic signaling downstream of Fas has been observed to be a regulator of T cell activation ^33–35^. To evaluate whether Fas-FasL signaling affects Th2 differentiation, we used CRISPR-Cas9 mediated gene targeting to ablate either *Fas* or *Fasl* in naïve CD4 T cells prior to Th2 polarization *in vitro* (Fig S6A, B). WT cells treated with *Fas* or *Fasl* targeting gRNAs showed no changes in IL-4, IL-13 or IFNγ production (Fig 6D, E). However, *Fas* or *Fasl* targeted *Pik3cd^E1020K/+^* cells showed significantly reduced percentages of both Th2 cytokine producing cells (IL-4, IL-13) and notably, IFNγ-expressing cells under Th2 conditions (Fig 6D, E). These results suggest that Fas-FasL signaling modulates aberrant cytokine production in *Pik3cd^E1020K/+^* Th2 cells.

We next evaluated the consequences of Fas-FasL ablation on *Pik3cd^E1020K/+^*cell signaling. Surprisingly, we observed significantly diminished pAKT(T308), a proximal readout of PI3Kδ activity, in *Fas* or *Fasl* targeted *Pik3cd^E1020K/+^* Th2 cells compared to control, with activation resembling control WT cells (Fig 6F, S6C). pFoxo1(S256) and pS6(S240/44) were similarly reduced by *Fas* or *Fasl* ablation in *Pik3cd^E1020K/+^*Th2 cells (Fig 6F, S6C), as was phosphorylation of p65/RelA (Fig S6D), a component of NFkB that has previously been found to be activated by non-canonical Fas signaling ^36–38^. These results suggest that Fas-FasL activity amplifies T cell signaling in *Pik3cd^E1020K/+^* cells.

To determine whether the effects we observed resulted from FasL stimulation of Fas, or signaling through the FasL intracellular domain ^39,40^, we used a recombinant FasL with an engineered leucine zipper domain that facilitates multimerization (FasL-LZ) to mimic effects of increased FasL. Both WT and *Pik3cd^E1020K/+^* Th2 cells differentiated in the presence of FasL-LZ showed significantly increased pS6(S240/44) compared to control Th2 cells (Fig 6G, H). This effect was dependent on the presence of Fas; Fas-deficient cells were completely resistant to the effects of FasL, whereas FasL-deficient cells retained the ability to respond to the inclusion of FasL-LZ in culture (Fig 6G, H). Identical trends were observed for pFoxo1(S256) and pAKT(T308), with FasL-LZ stimulation further exacerbating activation of *Pik3cd^E1020K/+^*Th2 cells (Fig 6H, S6E, F). Thus, engagement of Fas amplifies signaling pathways in *Pik3cd^E1020K/+^* CD4 T cells.

### Fas co-stimulates TCR signaling

To investigate mechanisms behind Fas-induced amplification of PI3K pathways, we fused the cytoplasmic tail of Fas to a biotin ligase (BioID2), which we expressed in a Jurkat cell line that lacks endogenous Fas (Fig 7A). BioID screening allows for the identification of potential protein-protein interactions through the transfer of biotin groups from a tagged bait protein to interacting partners, which can be isolated and identified by mass spectrometry. Using this approach, we analyzed 7077 total proteins, with FasL-LZ stimulated Fas-BioID cells showing specific enrichment of 170 proteins relative to Jurkat cells expressing BioID alone (Fig 7A, Supplementary Table 4). Among these Fas-BioID specific proteins, we identified known Fas associated proteins including Fas itself, FADD and Caspase 8; these latter two were only labeled upon FasL-LZ stimulation, confirming the fidelity of our Fas-BioID approach (Fig 7B, S7A). In addition to known Fas-associated proteins, we also observed significant interactions with multiple TCR signaling molecules including CD3e and Lck (Fig 7B, S7A). Pathway enrichment analysis of Fas-BioID+FasL-LZ specific proteins identified phosphorylation of CD3 and TCRz chains as the top enriched pathway (Fig 7C).

**Figure 7.**
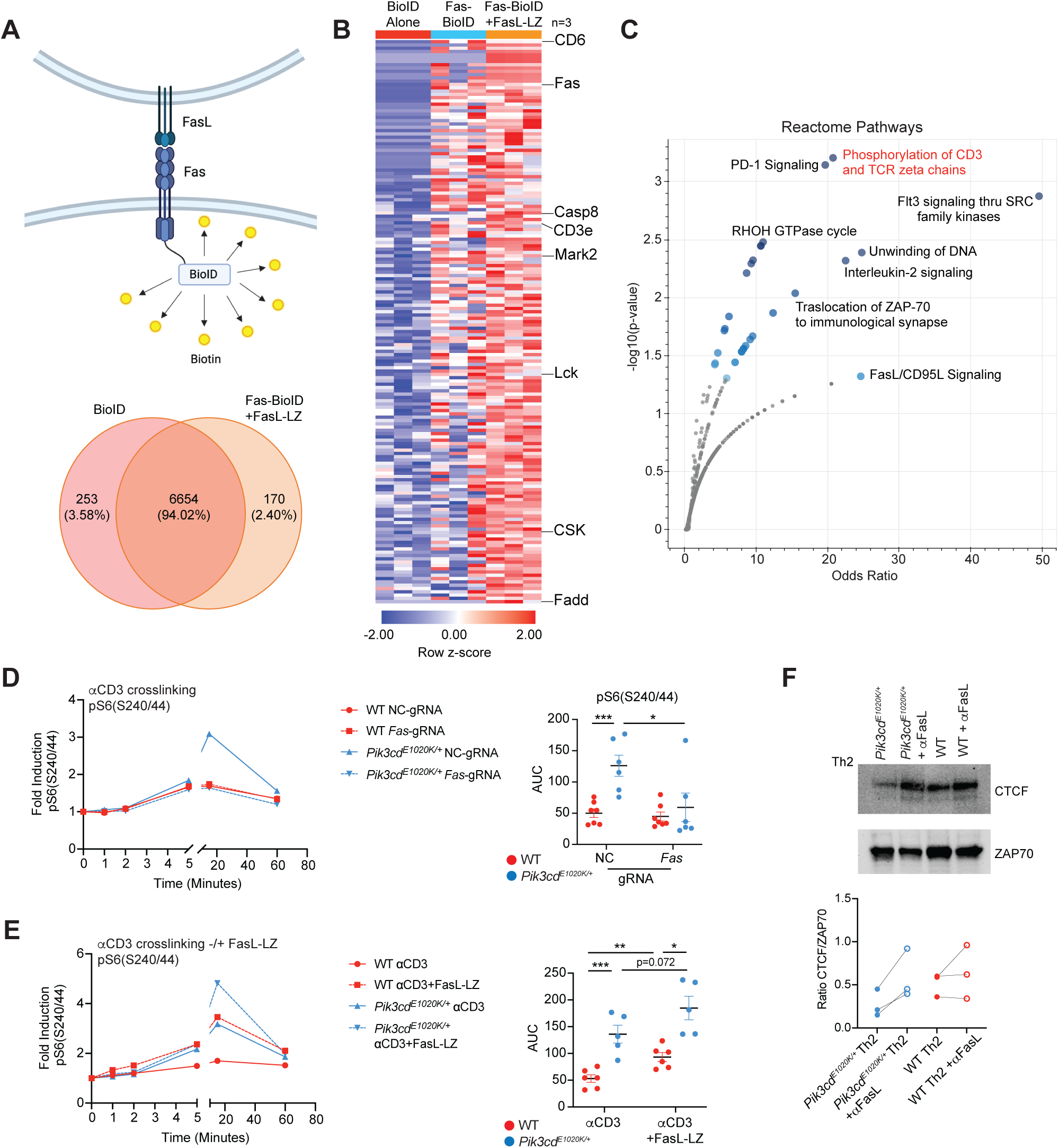
Fas interacts with the TCR complex and co-stimulates TCR signaling. A) Top, Fas was tagged with Bio-ID2 on its intracellular C-terminus (Fas-BioID) and stably expressed in Fas deficient Jurkat cells. Fas-BioID Jurkat cells were cultured in the presence or absence of recombinant multimeric FasL (FasL-LZ). As a control, BioID alone also was stably expressed in Jurkat cells. Bottom, Venn diagram showing proteins identified following mass spectrometry analysis of biotinylated proteins (streptavidin pulldown) that were common between or specific to BioID alone Jurkat cells versus Fas-BioD + FasL-LZ (p<0.05, n=3 for all groups). B) Heatmap (row z-score) showing % normalized spectral abundance of proteins specifically upregulated in Fas-BioID + FasL-LZ Jurkat cells relative to BioID alone Jurkat cells. C) Pathway enrichment of Reactome gene sets was performed using Enrichr ^73^, with significantly enriched gene sets colored in blue; proteins specifically upregulated in Fas-BioID + FasL-LZ Jurkat cells relative to BioID alone Jurkat cells were used as input for pathway enrichment. D) Naïve CD4 T cells from WT and *Pik3cd^E1020K/+^* mice were nucleofected with Cas9-gRNA complexes containing negative control or *Fas* targeting gRNAs and underwent aCD3/CD28 stimulation in the presence of hIL-2 for 72 hours. Cells were subsequently rested in serum free media, underwent aCD3 (1μg/mL) crosslinking over a time course (0, 1, 2, 5, 15, 60 minutes) and were analyzed by flow cytometry. n=6-7 for each group, 6-7 independent experiments. Left, fold induction of pS6(S240/44) over time for the indicated groups. Fold induction was calculated using pS6(S240/44) MFIs normalized to the 0 timepoint of the corresponding sample. Right, area under the curve (AUC) quantification of pS6(S240/44) time courses for the indicated groups. E) Naïve CD4 T cells from WT and *Pik3cd^E1020K/+^* mice underwent aCD3/CD28 stimulation in the presence of hIL-2 for 72 hours. Cells were subsequently rested in serum free media, underwent aCD3 (1μg/mL) crosslinking in the presence or absence of FasL-LZ over a time course (0, 1, 2, 5, 15, 60 minutes) and were analyzed by flow cytometry. n=6-7 for each group, 6-7 independent experiments. Left, fold induction of pS6(S240/44) over time for the indicated groups. Fold induction was calculated using pS6(S240/44) MFIs normalized to the 0 timepoint of the corresponding sample. Right, AUC quantification of pS6(S240/44) time courses for the indicated groups. F) Naïve CD4 T cells were Th2 polarized in the presence or absence of an aFasL blocking antibody. Top, Western blot showing CTCF and Zap70 protein in the indicated groups. Bottom, quantification of CTCF protein expressed as a ratio of CTCF/Zap70. n=3 for each group, 3 independent experiments. Statistical comparisons were made using ratio paired t-tests. *p<0.05, **p<0.01, ***p<0.001.

To evaluate whether Fas directly influences TCR signaling, we activated control and *Fas*-targeted CD4 T cells for 3 days, then rested cells in serum free media without cytokines, followed by acute aCD3 crosslinking. Using pS6(S240/44) as a readout of downstream TCR signaling, we confirmed significantly elevated phosphorylation in *Pik3cd^E1020K/+^* cells relative to WT cells (Fig 7D). However, Fas-deficient *Pik3cd^E1020K/+^* CD4 T cells displayed reduced pS6(S240/44) in in response to aCD3 crosslinking (Fig 7D). Additionally, Fas-deficient *Pik3cd^E1020K/+^*CD4 T cells also showed diminished pAKT(T308) and pERK(T202/04) responses compared to control cells (Fig S7B), confirming broad effects of the loss of Fas engagement on multiple T cell signaling pathways in activated PI3Kδ CD4 T cells.

Decreased TCR signaling was not observed in Fas-deficient WT CD4 T cells (Fig 7D, S7B), likely due to their low expression of FasL (Fig 6A). To probe Fas-mediated regulation of TCR signaling in WT cells, we again activated CD4 T cells for 3 days, rested cells in serum free media and acutely crosslinked CD3 in the presence or absence of recombinant FasL-LZ. Using pS6(S240/44) as a readout, we observed significantly elevated phosphorylation in WT CD4 T cells stimulated through the TCR in the presence of FasL-LZ compared to control TCR-stimulated cells (Fig 7E). Indeed, WT cells stimulated with aCD3 in the presence of FasL-LZ showed similar amplification of pS6(S240/44) as *Pik3cd^E1020K/+^*cells stimulated with aCD3 alone. Similarly, induction of pAKT(T308) and pERK(T202/04) was increased by FasL-LZ co-stimulation of WT cells (Fig S7C), again suggesting broad downstream effects of Fas engagement, consistent with the labeling of proximal TCR signaling components by BioID. Moreover, FasL-LZ co-stimulation further elevated pS6(S240/44), pAKT(T308) and pERK(T202/04) in *Pik3cd^E1020K/+^*CD4 cells (Fig 7E, S7C).

Given the ability of Fas/FasL signaling to reshape *Pik3cd^E1020K/+^* CD4 T cell signaling and differentiation, we examined the consequences of blocking FasL on CTCF expression. Using a FasL blocking antibody, we observed increased CTCF protein in αFasL-treated *Pik3cd^E1020K/+^*Th2 cells compared to control counterparts (Fig 7F). Therefore, signaling amplification via Fas-FasL regulates both T cell activation and CTCF protein expression in *Pik3cd^E1020K/+^* cells.

Together, these results suggest that activated PI3Kδ promotes altered gene expression and epigenetic modifications via a Foxo1-FasL-Fas amplification loop that induces IFNγ and Type I inflammatory gene expression, alters CTCF availability and disrupts Th2 lineage restriction.

## Discussion

We have uncovered a unique PI3K signaling circuit that enforces transcriptional and epigenetic programs regulating Th2 lineage specification. Activation of PI3Kδ downstream of TCR drives elevated IL-2 signaling, including increased production of IL-2 as well as exaggerated IL-2- induced Foxo1 inactivation that, in turn, further increases IL-2 expression, enforcing a positive amplification loop that drives IFNγ expression and prevents Th2 lineage restriction. We have further identified Foxo1-repressed Fas-FasL signaling as a novel mechanism affecting T cell activation, via amplification of signaling from the TCR. This amplification of PI3Kδ signaling facilitates chromatin reorganization, modulating the availability of CTCF, and is associated with profound changes in the cellular transcriptional and epigenetic landscape. Together, this dynamic signaling loop, generates a self-reinforcing signaling circuit that results in robust CD4 T cell activation and differentiation that likely contributes to PI3Kδ hyperactivation seen in APDS.

APDS patients show numerous pathologies in the respiratory tract, including frequent recurrent respiratory infections and bronchiectasis ^41^. Nonetheless, despite reports of increased incidence of asthma ^5,6^, it remains unclear whether patients show a classical eosinophilic asthma versus other lung pathology. *In vitro*, CD4 T cells from APDS patients and *Pik3cd^E1020K/+^* mice show increased IL-4 and IL-13 production under Th2 conditions ^5,26^, yet how this response proceeds in patients *in vivo* is unclear. In our mouse model, we found a highly dysregulated response to house dust mite in lung tissue as a consequence of PI3Kδ hyperactivation, with increased infiltration of CD4 T cells, yet these mice lack a Th2 response profile and show strong IFNγ signatures and increased neutrophils. Indeed, a high proportion of circulating Tfh-like cells from APDS patents express CXCR3, a marker of IFNγ responses ^42^; APDS patients also have elevated serum CXCL10, an IFNγ-induced chemokine ^43^. Furthermore, despite reports of Th2 pathologies, APDS patients do not show elevated serum IgE ^5,6^. These somewhat contradictory findings could be due to inappropriate production of IFNγ by CD4 T cells expressing activated PI3Kδ and strong polarizing effects of IFNγ as has been seen in mouse models of lung disease ^44^, although effects of activated PI3Kδ are likely to be context- and exposure-specific and the lack of IgE may result from immunoglobulin class-switching defects. It is of note that PI3Kδ inhibitors have been examined in certain lung diseases such as COPD; whether their use may be beneficial in the context of neutrophilic asthma may warrant closer examination ^45–47^.

The role of Foxo1 as a transcriptional repressor has been described in various cell types; our data now demonstrate the interplay between IL-2, PI3Kδ and Foxo1 in specifically shaping CD4 lineage decisions. In addition to uncovering a novel role of Foxo1 in repressing Th2 lineage cytokines IL-4 and IL-13, we further found that loss of Foxo1 mediates the induction of the Th1 program under Th2 inducing conditions, highlighting the requirement of balanced Foxo1 activity to ensure proper lineage determination. In addition, we show that Foxo1 repression results in elevated production of IL-2, driving further inactivation of Foxo1 and establishing a signaling circuit that allows for simultaneous adoption of multiple CD4 T cell lineages in *Pik3cd^E1020K/+^* cells. While the involvement of PI3K downstream of IL-2 signaling has been controversial ^12^, we see evidence for this amplification circuit in the context of activated PI3Kδ. Finally, an added consequence of Foxo1 inhibition by IL-2-PI3Kδ is the unleashing of *Fasl* expression, triggering a Fas-FasL signaling cascade that promotes exacerbated TCR signaling and a loss of Th2 lineage restriction.

Multiple PI3K-mediated pathways likely contribute to phenotypes associated with APDS, including activation of mTORC1 and downstream hypoxia-driven genes, altered metabolism, enhanced adhesion and increased senescence ^3,4,48^. Our data now suggest that non-apoptotic Fas-FasL signaling serves as a critical input exacerbating activation of *Pik3cd^E1020K/+^* CD4 T cells. Furthermore, the observation that Fas stimulation of WT CD4 T cells using recombinant FasL-LZ promoted T cell activation via amplification of PI3K and ERK signaling downstream of the TCR, suggests this circuit can be part of normal T cell activation. BioID screening revealed interactions between TCR components and Fas, providing a basis for the ability of Fas to potentiate TCR signaling. Indeed, previous work has also suggested cooperativity between Fas and TCR components regulating both proximal and distal TCR signaling events ^34,35^; other evidence points towards a role for Fas signaling promoting acquisition of effector phenotypes in T cells, suggesting important roles for these pathways in normal T cells ^33,49,50^. Intriguingly, Fas has also been shown to be critical for resolution of hypersensitivity and airway reactivity in a mouse model of allergic asthma, in part by driving expression of IFNγ ^50^. Given the propensity for Fas signaling to induce cell death in T cells, this mode of co-stimulation presents an interesting dichotomy. On one hand, strong signaling through Fas generally induces cell death; yet, what determines whether Fas induces T cell activation is less clear. Modulation of FasL expression levels, subcellular trafficking ^36^ or altered activation of downstream signaling components such as PI3K, or Src family kinases as has been seen in glioblastoma cell lines ^51^, may be key to harnessing this pathway, with context dependent regulation of FasL guiding Fas signaling outcomes that promote or inhibit immune responses. Furthermore, this amplification loop may have broad implications for other outcomes associated with PI3K activation where non-apoptotic Fas signaling has been shown to regulate a range of cellular processes in a variety of cell types ^36,50–57^.

Our findings of dysregulated epigenetic remodeling driven by hyperactivated PI3Kδ further highlight an unappreciated role of this signaling pathway in influencing chromatin organization during CD4 T cell differentiation. The increased accessibility of Th1 promoting motifs like Tbet, Runx and IRF, and a loss of Th2 associated motifs such as GATA3 and STAT6, in *Pik3cd^E1020K/+^* CD4 Th2 cells are consistent with the loss of Th2 lineage restriction we observed. However, more surprising was the attenuation of CTCF motif accessibility and CTCF-DNA binding in the context of PI3Kδ hyperactivation. The importance of CTCF in reshaping chromatin architecture in CD8 T cells has been highlighted by recent data describing dynamic changes in CTCF binding in the transition from naïve to effector and memory T cell differentiation states ^58–60^. In CD4 T cells, dynamic changes in CTCF activity occur near the *Ifng* and Th2 cytokine loci during Th1 and Th2 differentiation respectively ^22,61–64^. However, we found a more global pattern of impaired CTCF activity that corresponded to genomic regions associated with multiple transcriptional regulators, including Foxo1; this is likely to be a more universal feature of PI3K activity, as that repression of CTCF motif accessibility is also seen in *Pik3cd^E1020K/+^* CD8 T cells ^65^. It is therefore of interest that recent work indicates that decreased CTCF levels are associated with increased variability in gene expression ^66^. The overall profile of reduced CTCF protein in *Pik3cd^E1020K/+^* Th2 cells highlights the highly dysregulated nature of PI3Kδ hyperactivation, which leads to global changes in chromatin organization and loss of Th2 lineage restriction.

Integration of PI3Kδ signaling with the transcriptional and epigenetic networks that orchestrate CD4 T cell differentiation requires a delicate balance of signals to ensure both robust activation and lineage specification. Excessive signaling through PI3Kδ disrupts this balance, causing a downstream flood of cellular dysregulation at multiple levels. The ability of PI3Kδ to amplify its own activation through linking IL-2, Foxo1 and Fas-FasL signaling demonstrates a hard-wired mechanism for sustaining T cell activation during the acquisition of effector function. However, rewiring of this signaling circuit by PI3Kδ hyperactivation incites inappropriate amplification of signaling responses that cause T cell dysregulation and a loss of T cell lineage identity. How this contributes to phenotypes associated with altered PI3Kδ activity in immunodeficiencies and other diseases remains an important question.

## Methods

### Resource Availability

#### Lead Contact

Further information and requests for resources and reagents should be directed to the lead contact, Pamela L. Schwartzberg.

### Materials availability

No unique reagents were generated in this study. *Pik3cd^E1020K/+^* mice are available with a completed Materials Transfer Agreement.

### Data and code availability

This paper does not report original code.

Sequence data has been deposited under GSE214871, GSE278337, GSE278564 and GSE277881 will be available upon publication. Other data is included in supplemental materials.

Any additional information required to reanalyze the data reported in this paper is available from the lead contact upon request.

### Experimental Model Details

#### Mice

*Pik3cd^E1020K/+^* mice have been described previously and were backcrossed > 10 times to C57BL6/J. For all experiments, age and sex matched mice between 8-12 weeks of age were used. For adoptive transfer experiments, TCRa-deficient mice (B6.129S2-*Tcra^tm1Mom^*) were obtained from NIAID-Taconic contract facility and used as recipients. Mice were maintained and treated under specific pathogen free (SPF) conditions in accordance with the guidelines of NHGRI (protocol G98-3), NIAID (protocol LPD-6 and LISB-22E) and the Animal Care and Use committees at the NIH (Animal Welfare Assurance #A-4149-01).

### Method Details

#### Antibodies and flow cytometry

All flow cytometry antibodies were purchased from BD, Biolegend, Thermo Fisher or Cell Signaling Technologies. Surface antibody staining was performed in FACS buffer (PBS supplemented with 1% FBS and 1mM EDTA) for 30 minutes at 4°C. For transcription factor staining, cells were stained using the Foxp3 staining kit (Fixation for 25 minutes at room temperature, intracellular staining for a minimum of 1 hour at 4°C). For intracellular cytokine analysis, cells were restimulated with PMA (50ng/mL) and Ionomycin (500ng/mL) in the presence of GolgiStop (1/2000) at 37°C for 4 hours. Cytokine staining was performed using fixation in 4% PFA (25 minutes at room temperature) followed by intracellular staining in 0.5% Trition X-100, 0.1% BSA in PBS (minimum of 1 hour at 4°C). Staining of phospho-proteins was performed using fixation in 4% PFA (25 minutes at room temperature) followed by permeabilization using ice-cold 100% methanol (1 hour at 4°C) and intracellular staining in 0.5% Trition X-100, 0.1% BSA in PBS (minimum of 1 hour at 4°C). Cells were acquired using a Fortessa (BD) flow cytometer and analyzed using FlowJo software (v10.8.1)

#### House dust mite induced airway inflammation

House dust mite extract (*Dermatophagoides pteronyssinus)* was purchased from Greer laboratories and is extensively tested for endotoxin contamination. Mice were anesthetized with isoflurane and treated intranasally with 30μL of house dust mite extract over the course of 18 days. On days 0 and 7 mice were treated with 200μg of house dust mite extract. On days 14, 16 and 18 mice were treated with 50μg of house dust mite extract. Following the last sensitization, mice were euthanized on day 20 and lung tissue was harvested for analysis. Lung tissues were dissected and the left lung was used to make tissue homogenates for luminex analysis, post-caval lobe was fixed for histological analysis and superior, middle and inferior lobes were pooled for digestion and flow cytometric analysis. To prepare single cell suspensions for flow cytometry, lung tissue was cut into small pieces and digested in 0.25mg/mL of Liberase DL for 45 min at 37°C. Tissue was subsequently processed through a 70μm filter and treated with ACK lysis buffer to remove red blood cells. Single cell suspensions were reconstituted in complete media, counted and used for flow cytometry analysis. For adoptive transfer experiments, naïve CD4 T cells were isolated from spleen and lymph nodes of WT or *Pik3cd^E1020K/+^* mice using Miltenyi naïve CD4 T cell isolation kits and 1×10^6^ cells per mouse were injected intraperitoneally into TCRa KO mice. Following injection, mice were rested for 14 days before initiating the house dust mite protocol outlined above.

#### Histology

The post-caval lobe of the lung was fixed in 10% formalin (buffered, 7.2 pH) and embedded in paraffin. Embedded tissues were sectioned (5mm thick), mounted on glass slides and underwent Hematoxylin and Eosin (H&E) staining to examine immune cell profiles. All slides were digitized using an Aperio AT2 scanner (*Leica Biosystems, Wetzlar, Germany*).

The quantification of eosinophils was based on images from 10 randomly selected fields of lung sections per animal, captured at 40X magnification. All counts were carried out using *Aperio ImageScope* software (*Leica Biosystems, Wetzlar, Germany*). Eosinophils were identified by the bilobed shape of their nucleus and acidophilic cytoplasm and, using the cursor and program tools, they were marked and counted (results expressed in absolute numbers).

The assessment of inducible bronchus-associated lymphoid tissue (iBALT) formation in mouse lungs was carried out by morphometric analysis as previously described ^67^ and 10 images were captured at 10X magnification to cover the entire area of the histological lung section using *Aperio ImageScope* software (*Leica Biosystems, Wetzlar, Germany*). iBALT was identified and evaluated by a pathologist. The iBALT was identified by similar architecture to conventional secondary lymphoid organs, being found in the perivascular space surrounding large blood vessels and along the lung’s airways. The iBALT area was quantified using the QuPath v0.4.3 software ^68^ with results expressed as μm^2^.

The total mucosal area of the bronchi and bronchioles was also obtained by morphometric analysis. All bronchial and bronchial mucosal area was visualized by 10X magnification and captured using *Aperio ImageScope* software (*Leica Biosystems, Wetzlar, Germany*) and then were manually measured to obtain the total area in µm^2^, using QuPath v0.4.3 software. Subsequently, the same sections (analyzed previously) were visualized by the 4x magnification, captured using *Aperio ImageScope* software (*Leica Biosystems, Wetzlar, Germany*) and the length of each mucosa was manually measured in micrometers to obtain the actual proportion of the mucosal surface analyzed for each mouse. Then, the ratio of the total area (µm^2^) of mucosal of each animal was normalized to the smallest mucosal length to obtain a value representing the mucosal area (µm^2^) of each mouse.

#### Analysis of airway resistance

Mice were anesthetized by intraperitoneal injection of a ketamine (100mg/kg) and xylazine (20mg/kg) mixture. Mice subsequently underwent transtracheal intubation with a 20-gage SURFLO Teflon intravenous catheter (Santa Cruz Animal Health, Ft Worth, Tex). Mice were administered vecuronium bromide (1mg/kg) and mechanically ventilated using a FlexiVent respirator (SCIREQ). Respiratory system resistance (*R*_rs_) was measured after inhalation of PBS and increasing concentrations of methacholine.

#### Measurement of lung cytokines and chemokines in tissue homogenates

Following lung dissection, left lobes were isolated and added to an ice-cold mixture of 500μL RIPA buffer supplemented with protease inhibitor. Using Bertin hard tissue grinding tubes, samples were homogenized for 2 cycles of 20 seconds at 5500rpm in a Precellys Evolution (Bertin). Homogenates were subsequently centrifuged at 8000xg for 10 min to remove debris and supernatants were collected. Cytokine and chemokines were measured in homogenates using a 32plex Luminex kit (Millipore) and a Bio-Plex 200 analyzer (BioRad) per manufacturer’s instructions.

#### Measurement of house dust mite specific antibodies

Generation of HDM specific antibodies was determined by ELISA. Plates (Immunolon 4HBX) were coated with 5μg/mL HDM extract (Greer laboratories), incubated overnight at 4°C and blocked with 5% BSA in PBS. Samples were measured in duplicate with 50μL of serum samples added to HDM coated wells (IgG1(1/500), IgE(1/1), IgG2a(1/500), IgG3(1/500) and incubated at room temperature for 1 hour. Samples were washed with Tween/PBS and then probed with biotinylated rat anti-mouse IgG1, IgE IgG2a or IgG3 (1/250) and incubated at room temperature for 1 hour. Following washing, streptavidin conjugated HRP was added to samples and incubated at room temperature for 30 minutes. Samples were washed and treated with TMB (3, 3’,5,5’-tetramethylbenzidine), with 2N H_2_SO_4_ used to stop the reaction. Plates were analyzed using a multiplate reader (Molecular Devices) at absorbances of 450nm and 550nm.

#### Single cell RNAseq

Lungs were isolated from naïve and house dust mite (HDM) treated WT and *Pik3cd^E1020K/+^* animals (n=3 for all groups) and single cell suspensions were prepared as previously described. Live CD45^+^ cells were sorted using an Aria Fusion cell sorter (BD) and cells were labelled with unique hashtag antibodies (Biolegend). Labelled cells were pooled by group (WT naïve, *Pik3cd^E1020K/+^* naïve, WT HDM, *Pik3cd^E1020K/+^* HDM) with 1×10^4^ cells per donor (n=3 per group) used as input. Chromium Single Cell 3’ v2 libraries (10x Genomics) were generated following the standard manufacturers protocol and sequenced on one NovaSeq S2 and NovaSeq SP run at the Frederick National Laboratory for Cancer Research Sequencing Facility. The sequencing run was setup as a 28 cycles + 91 cycles non-symmetric run. All samples had sequencing yields of more than 206 million reads per sample. Samples were demultiplexed, allowing 1 mismatch in the barcodes. Over 93.7% of bases in the barcode regions had Q30 or above and at least 89.8% of bases in the RNA read had Q30 or above. More than 93.8% of bases in the UMI had Q30 or above. Data were analyzed using Cell Ranger v6.0.2 ^69^, following default parameters, mapping to the Mus musculus Genome Reference Consortium Mouse Build 38 (mm10, accession: GCA_000001635.2, https://www.ncbi.nlm.nih.gov/assembly/GCF_000001635.20/).

After alignment to mm10, followed by the quantification of RNA counts per cell, Seurat v4.1.1 ^70^ was used to filter the data and perform downstream analyses. Each dataset was visualized to confirm there were not bimodal distributions with an excess of cells with multiple hashtags and to determine the best cutoffs for cells lacking unique features or potential doublets. Cells were retained with more than 200 unique features but fewer than 3000, and with less than 5% of reads mapping to mitochondrial genes.

Following filtering, we integrated samples based on the variable features of each cell, and then clustered cells, following default clustering parameters, with a resolution of 0.25. Numbers of CD45^+^ lung cells identified for analysis are as follows: WT naïve (7077), *Pik3cd^E1020K/+^* naïve (5533), WT HDM (2111), *Pik3cd^E1020K/+^* HDM (14119). Cell types were identified in the integrated dataset using a combination of cluster specific gene expression data (Supplementary Table 1) and the ImmGen database in SingleR v1.8.1 ^71^. Seurat v4.1.1 was used to generate dimension plots showing clusters and cell types of these data, as well as to generate features plots of gene expression levels. To balance cell numbers for visualization in dimensional plots, data groups were downsampled to 2000 cells per condition. Enriched functional gene sets were identified using ShinyGo ^72^ and Enricher ^73^, with differentially expressed genes for indicated comparisons used as input.

To examine the response to interferon gamma (GO:0034341) gene set in all cell clusters identified, we used the AUcell R package ^74^. Briefly, we converted our data into an expression matrix of read counts per cell and identified whether genes contained in the response to interferon gamma gene set were expressed in each cell. We then used AUcell to generate an enrichment score of response to interferon gamma in all individual cells compared to all other genes. To visualize these results, we used Seurat to plot enrichment p-values as a dimension plot and barplot. We used Chi-squared tests to determine if there was a significant difference between WT and *Pik3cd^E1020K/+^*in the proportion of cells with highly significant (p<0.0005) enrichment p-values.

#### In vitro CD4 T cell polarization

Naïve CD4 T cells were isolated from spleen and lymph nodes of WT and *Pik3cd^E1020K/+^*mice using Miltenyi mouse naïve CD4 T cell isolation kits. Antigen presenting cells (APCs) were prepared by depleting T cells from splenocyte suspensions by labeling with CD4-FITC (1/200) and CD8a-FITC (1/200) and negatively selecting labelled cells using anti-FITC Microbeads.

Naïve T cells (2×10^5^) and APCs (1×10^6^) were mixed at 1:5 ratio and were plated into wells of a 48-well plate in 1mL of complete IMDM media (10% FBS, Penicillin/Streptomycin, Glutamine and 2-ME). Naïve CD4 T cells were activated with anti-CD3ε (1μg/mL) and anti-CD28 (3μg/mL) in the presence of cytokines and blocking antibodies. For Th1 conditions, cells were activated in presence of IL-12 (20ng/mL) and anti-IL-4 (10μg/mL). For Th2 conditions, cells were activated in the presence of IL-4 (40ng/mL) and anti-IL-12 (20μg/mL). Cells were cultured for 24-72 hours at 37°C (5% CO_2_) and analyzed at various timepoints during differentiation.

#### Bulk RNAseq

Live naïve CD4 T cells (CD4^+^CD8α^-^TCRβ^+^CD25^-^CD44^-^CD62L^+^) were sorted from spleens of WT and *Pik3cd^E1020K/+^* mice and were resuspended in 1mL of Trizol (n=3 for each group). Th1 and Th2 polarized cells (-/+ aIL2) were generated *in vitro* from WT and *Pik3cd^E1020K/+^* cells (as described) and live CD4^+^ cells were sorted directly into Trizol (1mL final volume). Total RNA was isolated from samples using phenol-chloroform extraction with GlycoBlue as a co-precipitant. RNA was quantified using Qubit RNA high sensitivity reagents and 30ng of total RNA was used as input for RNAseq library preparation. Libraries were prepared using the

NEBNext poly(A)+ mRNA magnetic isolation module and NEBNext Ultra RNA Library prep kit. Libraries were sequenced using the NovaSeq platform (Illumina) and underwent paired-end sequencing to produce between 262 and 540 million 100bp readpairs per sample, for a total of 13.8 billion read pairs.

Raw base calls were demultiplexed and converted into FastQ format with CASAVA 1.8.2. The raw data were processed using the OpenOmics/RNA-seek pipeline version 1.9.0 (https://github.com/OpenOmics/RNA-seek). Briefly, this pipeline assessed the quality of each sample using FastQC v0.11.9 ^75^, Preseq v2.0.3 ^76^, Picard tools v2.17.11 ^77^, FastQ Screen v0.9.3 ^78^, Kraken2 v2.0.8 ^79^, QualiMap ^80^, and RSeQC v2.6.4 ^81^. Adapters and low quality sequences were trimmed using Cutadapt v1.18 ^82^. The trimmed reads were aligned against the Mus musculus Genome Reference Consortium Mouse Build 38 (mm10, accession: GCA_000001635.2, https://www.ncbi.nlm.nih.gov/assembly/GCF_000001635.20/), using the splicing-aware aligner STAR version 2.7.6a ^83^ in per-sample two pass basic mode. Gene and transcript expression levels were estimated via RSEM v1.3.3 ^84^.

Transcripts were normalized and differentially expressed genes (DEG) called by quasi-likelihood F testing using *edgeR*. DEG call denotes >1.5 fold pairwise change and Benjamini-Hochberg (BH) adjusted *p* value < 0.05. Reads per kilobase per million (RPKM) were compiled with edgeR. An offset value of 1 was added to all RPKM and those failing to reach a value >2 RPKM in any genotype/condition were purged, as were micro-RNAs, sno-RNAs and sca-RNAs. RPKM+1 was the input for principal component analysis (*prcomp*).

*clusterprofiler* was used for hypergeometric testing (HGT) and geneset enrichment analysis (GSEA) against the KEGG, GO, Reactome and Molecular Signatures (MSigDB) databases or custom Foxo1 activated and repressed genesets (Supplementary Table 2). For HGT, input DEG lists were generated based on the above pairwise criteria. For GSEA, unabridged transcriptomes were ranked by Log2FC. GSEA plots were rendered with *enrichplot*, heatmaps with Morpheus (https://software.broadinstitute.org/morpheus) and all other plots with *ggplot2* or *Datagraph* (Visual Data Tools Inc., USA).

#### CRISPR-Cas9 gene targeting

Pre-designed CRISPR RNAs (crRNAs) were purchased from IDT and were reconstituted at 100μM (See table for sequence details). Trans-activating CRISPR RNA (tracrRNA) labelled with atto550 was purchased from IDT and reconstituted at 100μM. Guide RNAs (gRNAs) were generated by mixing crRNA and tracrRNA at a 1:1 ratio and duplexed by warming to 95°C for 5 minutes and were cooled at room temperature for 10 minutes before use. For each gene targeted, a mixture of 3 gRNAs was used to ensure efficient gene ablation. Ribonucleoprotein (RNP) complexes were formed by incubating gRNAs (3x1μL) with 1μL recombinant Cas9 (IDT) at room temperature for 5 minutes. Naïve CD4 T cells were isolated (as previously described), resuspended in 20μL Lonza P3 Nucleofection buffer and mixed with RNP complexes. Naïve CD4 T cells underwent nucleofection using P3 Primary Cell 4D-Nucleofector X Kit S cuvettes and the DN100 program of a Lonza 4D-nucleofector device. Following nucleofection, cells were washed and cultured in complete IMDM media supplemented with 5ng/mL IL-7. The IL-7 culture period ranged from 1-7 days, depending on the target gene of interest. Negative control crRNA #1 was purchased from IDT and used in all negative control conditions for CRISPR-Cas9 experiments.

#### Retroviral transduction

For retrovirus production, HEK 293T cells were co-transfected with GFP-pMIGR, Foxo1-WT-pMIGR or Foxo1-AAA-pMX (2μg) along with pCL-Eco helper plasmid (1μg). 48 hours post-transfection, cell supernatants were collected and incubated with 5x PEG-IT reagent (SBI) overnight at 4°C. Precipitated retrovirus was centrifuged at 1500xg for 30 minutes and supernatants were discarded. Retrovirus was centrifugated again at 1500xg for 5 minutes, supernatant discarded and concentrated retrovirus was reconstituted in complete IMDM. For retroviral transduction, naïve CD4 T cells were activated as described above. 18 hours post activation concentrated retrovirus (20μL) was mixed with polybrene A (8µg/mL final), added to 1mL of Th2 polarized cells in a 48 well plate and spinfected at 2000rpm for 1 hour at 37°C). Transduced cells were subsequently cultured at 37°C for a total of 72 hours, without changing differentiation media. Transduced cells were detected by flow cytometry using GFP expression.

#### Recombinant FasL-LZ

Recombinant FasL-LZ was engineered by fusing the extracellular domain of Fas ligand to a FLAG tag and an isoleucine zipper motif that facilitates self-oligomerization of the construct ^33^. This construct was transfected into HEK 293T cells and cell supernatants were collected. Recombinant FasL-LZ was purified using magnetic beads conjugated to anti-FLAG (Sigma). Quantification of recombinant protein was determined by ELISA (R&D). For all mouse CD4 T cell experiments, FasL-LZ was used at a concentration of 25ng/mL. For Jurkat BioID screening, FasL-LZ stimulation was performed at 10ng/mL.

#### Fas-BioID screening

Human *Fas* coding sequencing tagged with BioID2 (Addgene #80899) and P2A-EGFP at the C terminus was subcloned into the pMy retroviral backbone (Addgene # 163361) using Gibson assembly. Retrovirus was packaged using Plat-A cells (Cell Biolabs # RV-102). Fas deficient RapoC2 Jurkat cells were previously described ^85^. RapoC2 Jurkat cells were transduced with retrovirus expressing Fas-BioID or BioID alone by spinfection at 1500 x g at 32°C for 1 hour with 8 µg/mL polybrene (Sigma # TR-1003-G). Jurkat cells expressing Fas-BioID or BioID alone were enriched by FACS sorting using EGFP expression. BioID alone and Fas-BioID Jurkat cell lines were cultured in complete RPMI at 37°C (5% CO_2_), with 50μM Biotin (Sigma # B4501) added for the final 20 hours of culture. In addition, Fas-BioID Jurkat cells were treated with 10ng/mL FasL-LZ for the final 4 hours of culture. A minimum of 30×10^6^ cells were harvested for BioID only, Fas-BioID and Fas-BioID+FasL-LZ conditions, with 3 replicates prepared per group. Cell pellets were snap frozen prior to sample preparation. Cells were lysed using RIPA buffer (Sigma) with Halt inhibitor (Thermo) for 20 min on ice. Biotinylated proteins were pulled-down using streptavidin agarose beads (Pierce) at 4°C overnight. Mass spectrometry and data analyss were performed by Poochon Biotech (Frederick, MD). The LC/MS/MS analysis of tryptic peptides for each sample was performed sequentially with a blank run between two sample runs using a Thermo Scientific Orbitrap Exploris 240 Mass Spectrometer and a Thermo Dionex UltiMate 3000 RSLCnano System (Poochon Biotech). Peptides from trypsin digestion were loaded onto a peptide trap cartridge at a flow rate of 5 μL/min. The trapped peptides were eluted onto a reversed-phase Easy-Spray Column PepMap RSLC, C18, 2 μM, 100A, 75 μm × 250 mm (Thermo Scientific) using a linear gradient of acetonitrile (3-36%) in 0.1% formic acid. The elution duration was 110 min at a flow rate of 0.3 μl/min. Eluted peptides from the Easy-Spray column were ionized and sprayed into the mass spectrometer, using a Nano Easy-Spray Ion Source (Thermo) under the following settings: spray voltage, 1.6 kV, Capillary temperature, 275°C. Raw data files were searched against human protein sequences database using the Proteome Discoverer 2.4 software (Thermo, San Jose, CA) based on the SEQUEST algorithm. Carbamidomethylation (+57.021 Da) of cysteines was set as fixed modification, and Oxidation / +15.995 Da (M) and Deamidated / +0.984 Da (N, Q) were set as dynamic modifications. The minimum peptide length was specified to be five amino acids. The precursor mass tolerance was set to 15 ppm, whereas fragment mass tolerance was set to 0.05 Da. The maximum false peptide discovery rate was specified as 0.01.

#### Anti-CD3 crosslinking

For signaling studies with mouse CD4 T cells, naïve CD4 T cells were stimulated using APCs with anti-CD3ε (1μg/mL) and anti-CD28 (3μg/mL) in the presence of human IL-2 (50 IU/mL) for 72 hours at 37°C, as previously described. Following culture, activated CD4 T cells were purified using a Miltenyi mouse CD4 T cell isolation kit. Purified activated CD4 T cells were resuspended in serum-free IMDM media and rested at 37°C for 4 hours. Cells were subsequently counted and coated with anti-CD3ε (1μg/mL) in PBS for 15 minutes at 4°C. Cells were washed, resuspended in IMDM alone at 10×10^6^ cells/mL and incubated at 37°C for 10 minutes. Samples were subsequently stimulated by adding pre-warmed anti-armenian hamster (5μg/mL final) in IMDM to pre-incubated cells and cells were fixed at desired time points in PFA (4% final, 20 minutes at room temperature). For samples that underwent anti-CD3 + FasL-LZ stimulation, FasL-LZ (25ng/mL final) was mixed with anti-armenian hamster and added to cells in tandem with the initiation of TCR stimulation. Cells were stained following the phospho-staining protocol outlined above.

#### ATACseq

1×10^4^ viable *in vitro* polarized Th2 cells were sorted into 500μL of FACS buffer using an Aria Fusion Sorter (BD). Cells were pelleted and resuspended in 50uL of transposase mixture including 25μL 2xTD buffer (Illumina), 2.5μL TDE1 (Illumina), 0.5μL 1% digitonin (Promega) and 22uL water. Tagmentation was performed by incubation at 37°C for 30 minutes at 300rpm. Following incubation, DNA was purified using a Qiagen MinElute kit, eluting samples in 10uL. Purified tagemented DNA was PCR amplified using previously described primers ^86^, with 12 cycles of amplification. Amplified libraries were purified using a Qiagen PCR cleanup kit and sequenced for 50 cycles (paired-end reads) on a NovaSeq 6000 (Illumina). ATAC-seq was done in three biological replicates per genotype. Reads were mapped to the mouse genome (mm10 assembly) using Bowtie 0.12.8. In all cases, redundant reads were removed using FastUniq, and customized Python scripts were used to calculate the fragment length of each pair of uniquely mapped paired-end (PE) reads. Reads whose fragment lengths were less than 175 bp were kept and only one mapped read per unique genomic region was used to call peaks. Regions of open chromatin were identified by MACS (version 1.4.2) using a p-value threshold of 1 × 10−5. Only regions called in three replicates were used for downstream analysis. Peak annotation and motif analysis were performed with the Hypergeometric Optimization of Motif EnRichment program (HOMER) version 4.11.1 using the following parameter settings; “annotatePeaks.pl peak_file mm10 -size 1000 -hist 40 -ghist” and “findMotifsGenome.pl peak_file mm10 motif_folder -size given -preparsedDir tmp 2 > out”. All peaks detected in WT and *Pik3cd^E1020K/+^*Th2 were quantified for individual samples as follow; ‘annotatePeaks.pl peakfile mm10 -size given -raw -d ATAC-seq data’, and the differentially regulated peaks were called with a cut off value of FDR < 0.05 and log2FC > 0.5 using the following command; ‘getDiffExpression.pl rawcount.txt -AvsA -DESeq2 -fdr 0.05 -log2fold 0.5 -simpleNorm’. Downstream analyses and graph generation were performed with R 4.1.1.

#### CTCF CUT & Tag

Naïve (liveTCRb^+^CD4^+^CD8^-^CD44^-^CD62L^+^CD25^-^), *in vitro* Th1 polarized (liveCD4^+^) and Th2 polarized (liveCD4^+^) CD4 T cells from WT and *Pik3cd^E1020K/+^* mice were sorted using an Aria Fusion Sorter (BD), as previously described and then subjected to CUT&Tag using the CUT&Tag/v3 protocol (dx.doi.org**<u>/10.17504/protocols.io.bcuhiwt6</u>**) ^87^ as indicated on isolated nuclei from naïve CD4 T cells and *in vitro* polarized Th1 and Th2 cells from WT and *Pik3cd^E1020K/+^* mice. CUT&Tag utilized primary antibodies against CTCF (D31H2, Cell Signaling) and Rabbit mAB IgG XP Isotype Control (DA1E, Cell Signaling). All primary antibodies were diluted 1:50 with the corresponding guinea-pig anti-rabbit secondary diluted 1:100 accordingly. pA-Tn5 was sourced pre-loaded (C01070001, Diagenode). Primary antibody incubation occurred for 1 hr at RT with nutation and all other protocol steps as indicated in the native CUT&Tag protocol. Prepared libraries were normalized and pooled and prepared for sequencing via NovaSeq SP (Illumina) at the NHLBI Genomics Core. Raw reads were processed using CutRunTools/20200629 ^88^ with the mm10 mouse genome as reference. CTCF CUT&Tag signal data was RPKM normalized using deepTools/3.5.5 with extendReads enabled. Peak calling for CTCF was performed using SEACR/v1.3 with stringent settings and all fragment sizes incorporated. Peaks were further filtered by disassociation of peak summits belonging to the mm10 blacklist (CutRunTools) using bedtools/2.30.0 intersect. Peak signals were recalculated using deepTools computeMatrix +/- 200bp and further filtered by at least 2-fold signal change over cell-type summed IgG signal. Differential CTCF peaks from either genotype were at least +/- 4-fold change in signal between respective genotypes. CTCF motif scoring was evaluated by HOMER/4.11.1 findMotifsGenome.pl with option -size 200. *Pik3cd^E1020K/+^* induced and repressed CTCF peaks were visualized using deepTools and mapped to nearest gene using HOMER annotatePeaks default settings.

#### Western Blot

Naïve CD4 T cells were Th2 polarized as previously described and live CD4 T cells were enriched using Miltenyi CD4 T cell isolation kits. For each condition, 1×10^6^ cells were pelleted and cells were lysed using 50μL of 1% SDS in PBS (supplemented with protease inhibitor minitab (Sigma) and sodium orthovanadate) and 450mL 1% TritonX in PBS (containing inhibitors). Cell lysates were sheared using a 25-gauge needle with a 1cc syringe. Lysates were centrifuged at 14 000 rpm for 15 min (4°C). Lysates were prepared with reducing sample buffer and proteins were separated by SDS-PAGE and transferred to nitrocellulose. Membranes underwent blocking (TBS plus 5% BSA, 0.1% Tween-20) and were incubated with primary antibodies overnight (4°C). Subsequently, membranes were incubated with HRP-conjugated secondary antibodies for 60 minutes (room temperature). Signals were detected by chemiluminescence using a BioRad ChemiDoc Imaging System.

#### Statistical analysis

Data were analyzed using Prism v9 (GraphPad software). All graphs show mean -/+ SEM with *p<0.05, **p<0.01, ***p<0.001, ****p<0.0001. For unpaired comparisons, data were analyzed using unpaired t-tests. For paired comparisons, data were analyzed using ratio paired t-tests. For anti-CD3 crosslinking experiments, area under the curve (AUC) was calculated for individual signaling curves (baseline=1) and peak areas were compared using ratio paired t-tests.

## Acknowledgements

This research was supported in part by the Division of Intramural Research of NIAID, NIDDK and NIAMS, NIH. The authors would like to thank Jinfang Zhu and Andrea Pichler for discussion of the manuscript. The authors thank Kan Jiang and Hong-Wei Sun, Biodata Mining and Discovery Section at NIAMS and the NIAMS Sequencing Core Facility (Stephania Dell’Orso and Faiza Naz) for support with ATACseq. The authors additionally thank the NCI Center for Cancer Research Sequencing Facility for support with scRNAseq and bulk RNAseq experiments. AVV is funded by University of Miami, Department of Microbiology and Immunology start-up grant PG013596; University of Miami, Sylvester Comprehensive Cancer Center start-up grant PG012707.

## Supplemental Figure Legends

**Supplemental Figure 1.**
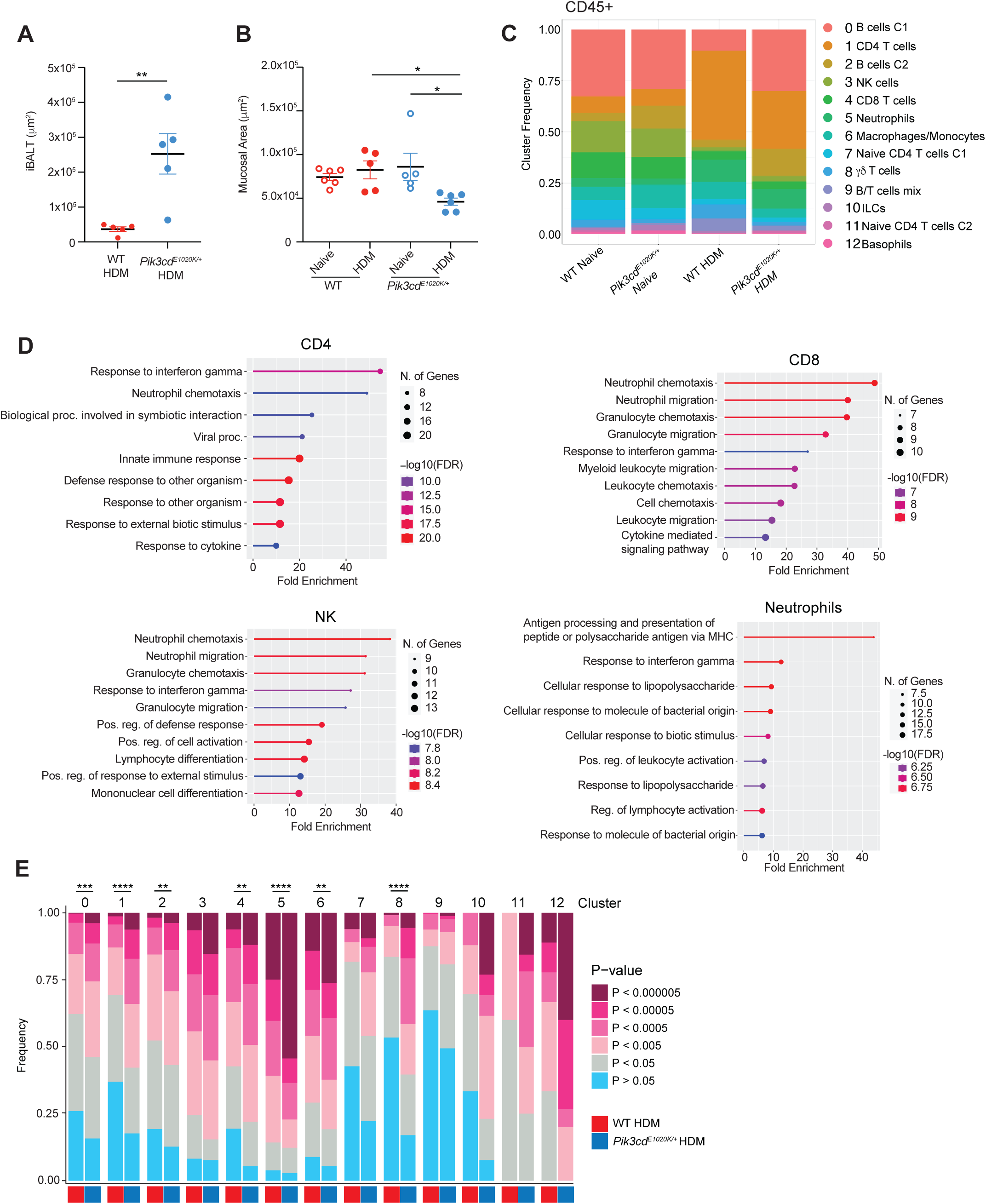
Altered responses to HDM in *Pik3cd^E1020K/+^* mice. HDM-sensitized mice from Fig. 1. A) Inducible bronchus associated lymphoid tissue (iBALT) area (μm^2^) measured according to H&E stained lung sections. B) Mucosal area (μm^2^) measured according to H&E stained lung sections. C) Frequencies of the indicated lung CD45^+^ immune cell populations (clusters 0-12) identified by scRNA-seq from naïve WT, naïve *Pik3cd^E1020K/+^*, HDM treated WT and HDM treated *Pik3cd^E1020K/+^* animals. D) Pathway enrichment analysis was performed using genes upregulated in HDM treated *Pik3cd^E1020K/+^* cells relative to HDM treated WT counterparts for the indicated cell types using scRNA-seq data of lung CD45^+^ immune cells. Pathway enrichment was performed using ShinyGo ^90^. E) Cluster specific analysis of frequencies of cells with the indicated enrichment p-value for the response to interferon-gamma gene set. Statistical comparison of WT and *Pik3cd^E1020K/+^* HDM treated groups for each cluster was performed (Chi-squared test), comparing frequencies of cells with p<0.0005. Unless otherwise indicated, statistical comparisons were made using unpaired t-tests. *p<0.05, **p<0.01, ***p<0.001, ****p<0.0001.

**Supplemental Figure 2.**
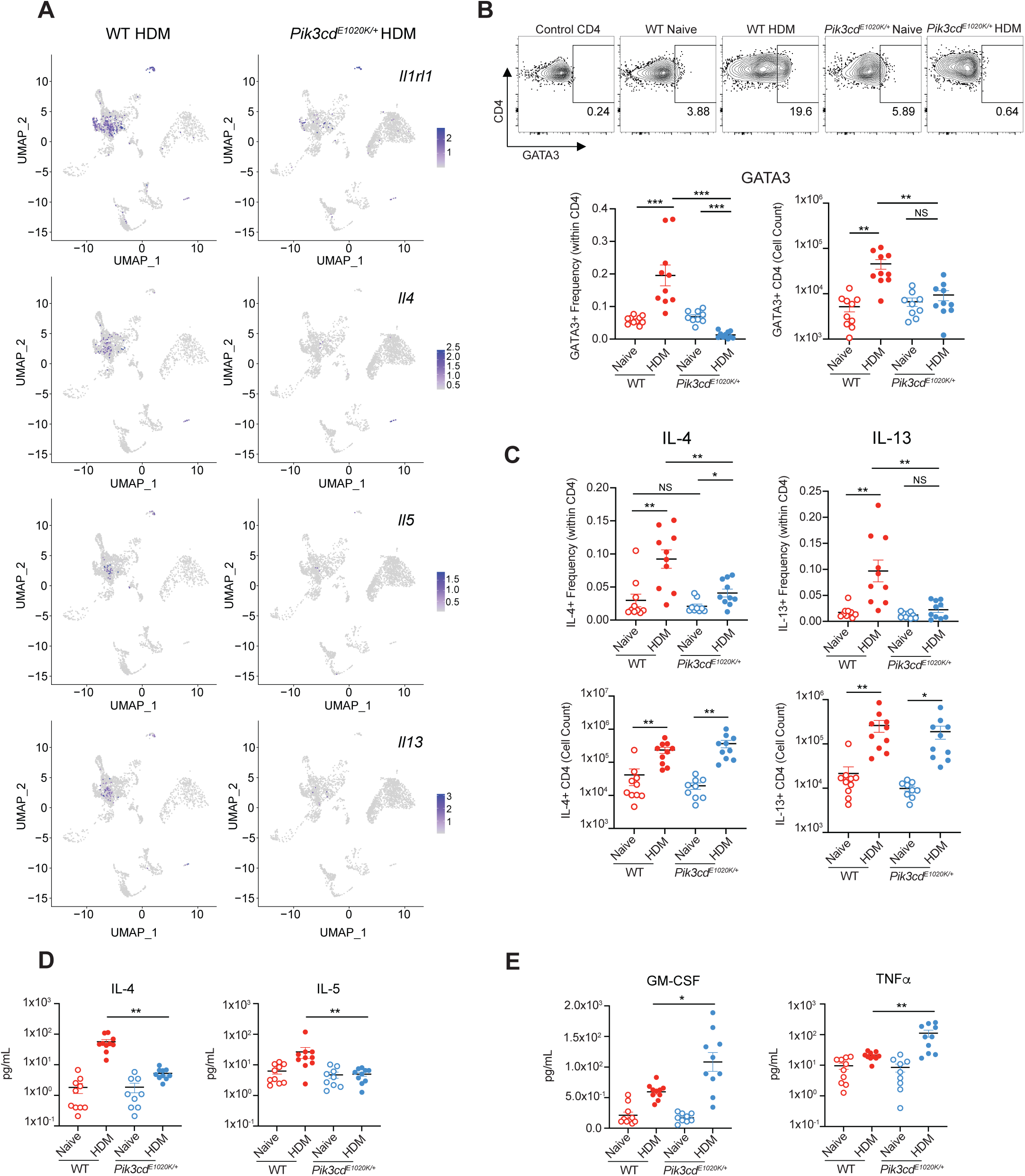
A) Feature plots showing expression of Th2 defining genes *Il1rl1*, *Il4*, *IL5* and *Il13* in scRNA-seq analyses of lung CD45^+^ immune cells from the indicated mice. B) Flow cytometry of lung CD4 T cells (liveCD45^+^TCRβ^+^CD4^+^CD8^-^) measuring GATA3 expression in the indicated populations. n=9-10 for each group, pooled from 2 independent experiments. Top, representative plots showing GATA3 and CD4 expression. Bottom, frequencies (left) and cell counts (right) of GATA3^+^ CD4 T cells in the indicated groups. C) Frequencies (top) and cell counts (bottom) of IL-4^+^ (left) and IL-13^+^ (right) lung CD4 T cells (liveCD45^+^TCRβ^+^CD4^+^CD8^-^CD4^+^) in the indicated groups, measured by intracellular staining and flow cytometry. n=9-10 for each group, pooled from 2 independent experiments. D, E) Cytokine concentrations (pg/mL) measured from lung homogenates from the indicated mice by Luminex. Th2 cytokines IL-4 and IL-5 are shown in D), Th1 cytokines GM-CSF and TNFα are shown in E). n=9-10 for each group, pooled from 2 independent experiments. Statistical comparisons were made using unpaired t-tests. *p<0.05, **p<0.01, ***p<0.001.

**Supplemental Figure 3.**
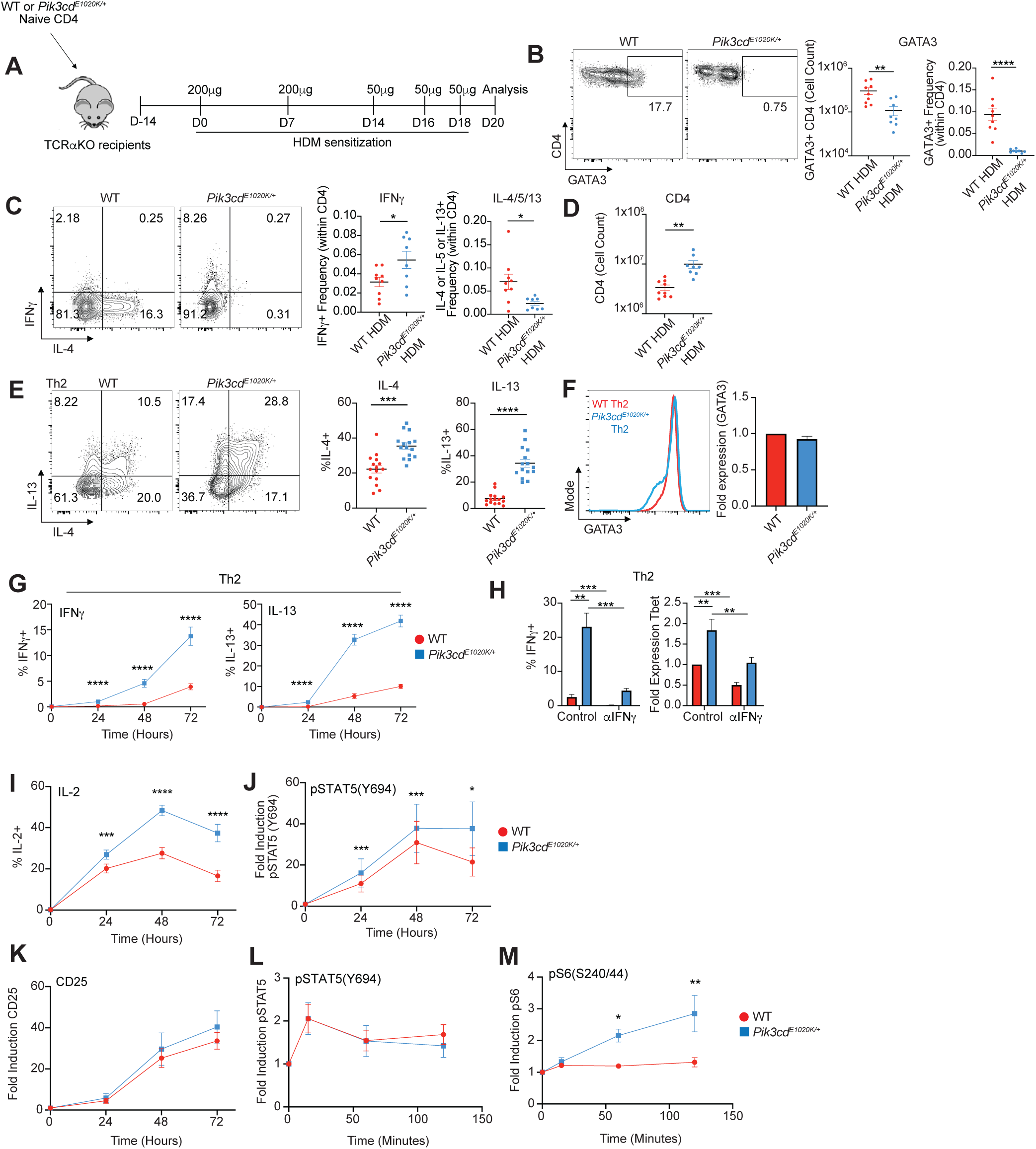
Cell-intrinsic alterations in Th2 differentiation. A-C) TCRa deficient recipient mice were injected with 1×10^6^ WT or *Pik3cd^E1020K/+^* naïve CD4 T cells 14 days prior to house dust mite (HDM) sensitization. HDM was administered intranasally as in Figure 1. n=9-10 for each group, pooled from 2 independent experiments. A) Experimental design. B) Left, Representative flow cytometry plots showing GATA3 and CD4 expression in lung CD4 T cells (live CD45^+^TCRβ^+^CD4^+^CD8^-^) from recipient animals receiving naive CD4 T cells from the indicated mice. Right, Cell counts (left) and frequencies (right) of GATA3^+^ CD4 T cells from the indicated groups. C) Left, Representative flow cytometry plots showing IFNγ and IL-4 expression in lung CD4 T cells (live CD45^+^TCRβ^+^CD4^+^CD8^-^) from recipient animals receiving naive CD4 T cells from the indicated mice. Right, Frequencies of IFNγ^+^ (left) and IL-4/5/13^+^ (right) lung CD4 T cells from the indicated groups. Frequencies of IL-4/5/13^+^ cells were calculated using Boolean (or) gating. D) Cell counts of lung CD4 T cells (live CD45^+^TCRβ^+^CD4^+^CD8^-^) from the indicated groups. E-M) Analyses of *in vitro* polarized Th2 cells. E) Left, Representative flow cytometry plots showing IL-4 and IL-13 staining in Th2 polarized live CD4^+^ cells. Right, Percentages of IL-4^+^ and IL-13^+^ cells from the indicated mice. n=15 for each group, 15 independent experiments. F) Left, representative flow cytometry histograms showing GATA3 expression in Th2 polarized live CD4^+^ cells from the indicated mice. Right, Fold GATA3 expression (MFI normalized to WT) in Th2 polarized live CD4^+^cells from the indicated mice. n=14 for each group, 14 independent experiments. G) Percentages of IFNγ^+^ (left) and IL-13^+^ (right) cells over a time course of Th2 differentiation (0, 24, 48 and 72 hours). n=10 for each group, 10 independent experiments. H) WT and *Pik3cd^E1020K^*^/+^ naïve CD4 T cells were Th2 polarized in the presence or absence of an αIFNγ blocking antibody. Left, percentages of IFNγ^+^ cells from the indicated groups. Right, fold Tbet expression (MFI normalized to control WT) in live CD4^+^ cells from the indicated groups. n=6-8 for each group, 6-8 independent experiments. I-K) Time course analysis (0, 24, 48, 72 hours) of IL-2 production, pSTAT5(Y694) and CD25 during Th2 polarization, measured by flow cytometry. n=7-10 for each group, 7-10 independent experiments. I) Percentages of IL-2^+^ cells over time from the indicated mice. J) Fold induction of pSTAT5(Y694) over time from the indicated mice. Fold induction was calculated using pSTAT5(Y694) MFIs normalized to the 0 timepoint of the corresponding genotype. K) Fold induction of CD25 expression from the indicated mice. Fold induction was calculated using CD25 MFIs normalized to the 0 timepoint of the corresponding genotype. L-M) Th2 polarized CD4 T cells from WT and *Pik3cd^E1020K/+^* were rested in serum free media for 4 hours and subsequently stimulated with hIL-2 over the indicated time course (0, 15, 60, 120 minutes). n=4-5 for each group, 4-5 independent experiments. L) Fold induction of pSTAT5(Y694) over time from the indicated mice, measured by flow cytometry. M) Fold induction of pS6(S240/44) over time from the indicated mice, measured by flow cytometry. Fold induction was calculated using MFIs (pSTAT5(Y694) or pS6(S240/44)) normalized to the 0 time point of the corresponding genotype. Statistical comparisons were made using unpaired t-tests (B-D) and ratio paired t-tests (E-M). *p<0.05, **p<0.01, ***p<0.001, ****p<0.0001.

**Supplemental Figure 4.**
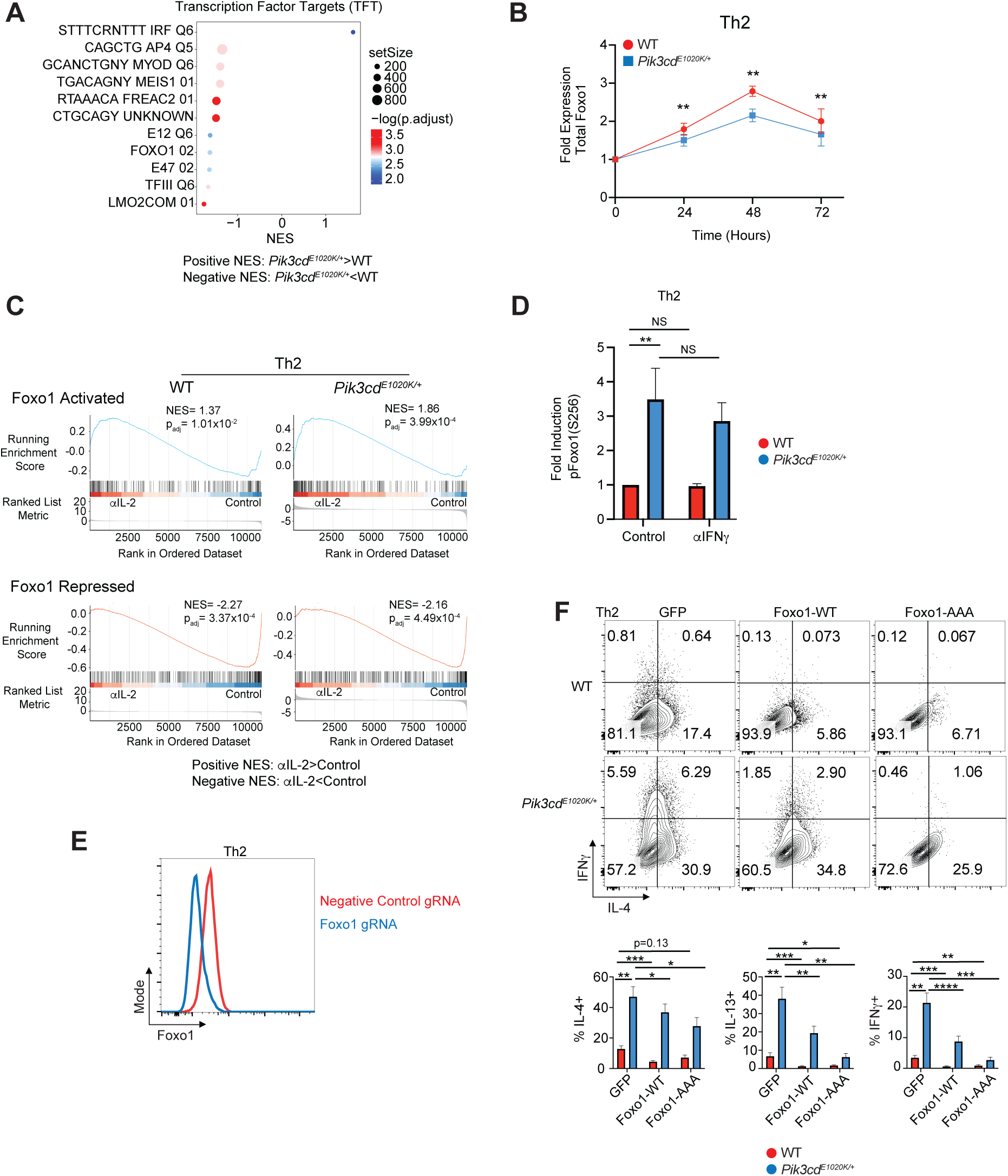
A) Gene set enrichment analysis (GSEA) comparing WT and *Pik3cd^E1020K/+^* transcriptomes for expression of transcription factor target (TFT) gene sets. B) Fold induction of Foxo1 expression in Th2 polarized live CD4^+^ cells, measured by flow cytometry and calculated by normalizing Foxo1 MFIs to the 0 time point of the corresponding genotype. n=5 for each group, 5 independent experiments. C) Gene set enrichment analysis (GSEA) comparing control and αIL-2 treated Th2 polarized CD4 T cell transcriptomes for the expression of Foxo1 activated (top) and repressed (bottom) gene sets in the indicated groups. D) Naïve CD4 T cells from the indicated mice were Th2 polarized in the presence or absence of αIFNγ blocking antibody and pFoxo1(S256) was measured in live CD4^+^ cells from the indicated groups by flow cytometry. Fold induction was calculated by normalizing pFoxo1(S256) MFIs to WT control cells. n=5 for each group, 5 independent experiments. E) Naïve CD4 T cells were nucleofected with Cas9-gRNA complexes containing negative control or *Foxo1* targeting gRNAs and underwent Th2 polarization. Representative flow cytometry histogram showing Foxo1 expression in cells from the indicated mice. F) Naïve CD4 T cells from WT and *Pik3cd^E1020K/+^* mice were transduced with GFP only, Foxo1-WT or Foxo1-AAA encoding retroviruses under Th2 polarizing conditions. Top, Representative flow cytometry plots showing IFNγ and IL-4 expression in live CD4^+^ T cells cultured under Th2 polarizing conditions from the indicated groups. Bottom, percentages of IL-4^+^ (left) and IFNγ^+^ (right) Th2 polarized liveCD4 T cells from the indicated groups. n=6 for each group, 6 independent experiments. Statistical comparisons were made using ratio paired t-tests. *p<0.05, **p<0.01, ***p<0.001, ****p<0.0001.

**Supplemental Figure 5.**
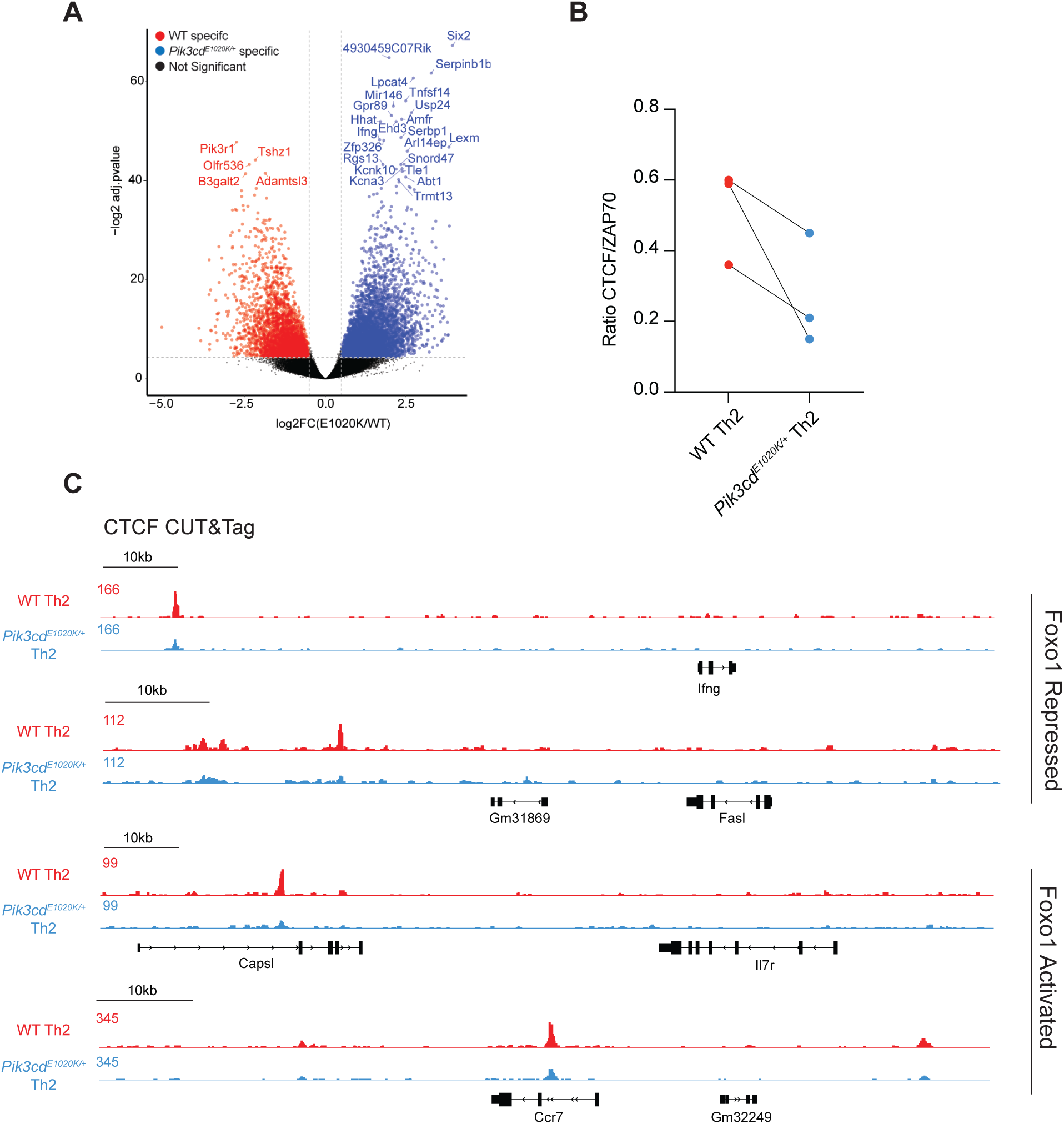
A) Naïve CD4 T cells from WT and *Pik3cd^E1020K/+^*mice underwent Th2 polarization and were examined by ATACseq. Volcano plot showing WT specific peaks in red and *Pik3cd^E1020K/+^* specific peaks in blue (fold change>1.5, p<0.05) B) Quantification of CTCF protein expressed as a ratio of CTCF/Zap70. Supporting data for Figure 5E. n=3 for each groups, 3 independent experiments. C) CTCF CUT&Tag tracks of *Ifng*, *Fasl*, *Il7r* and *Ccr7* loci.

**Supplemental Figure 6.**
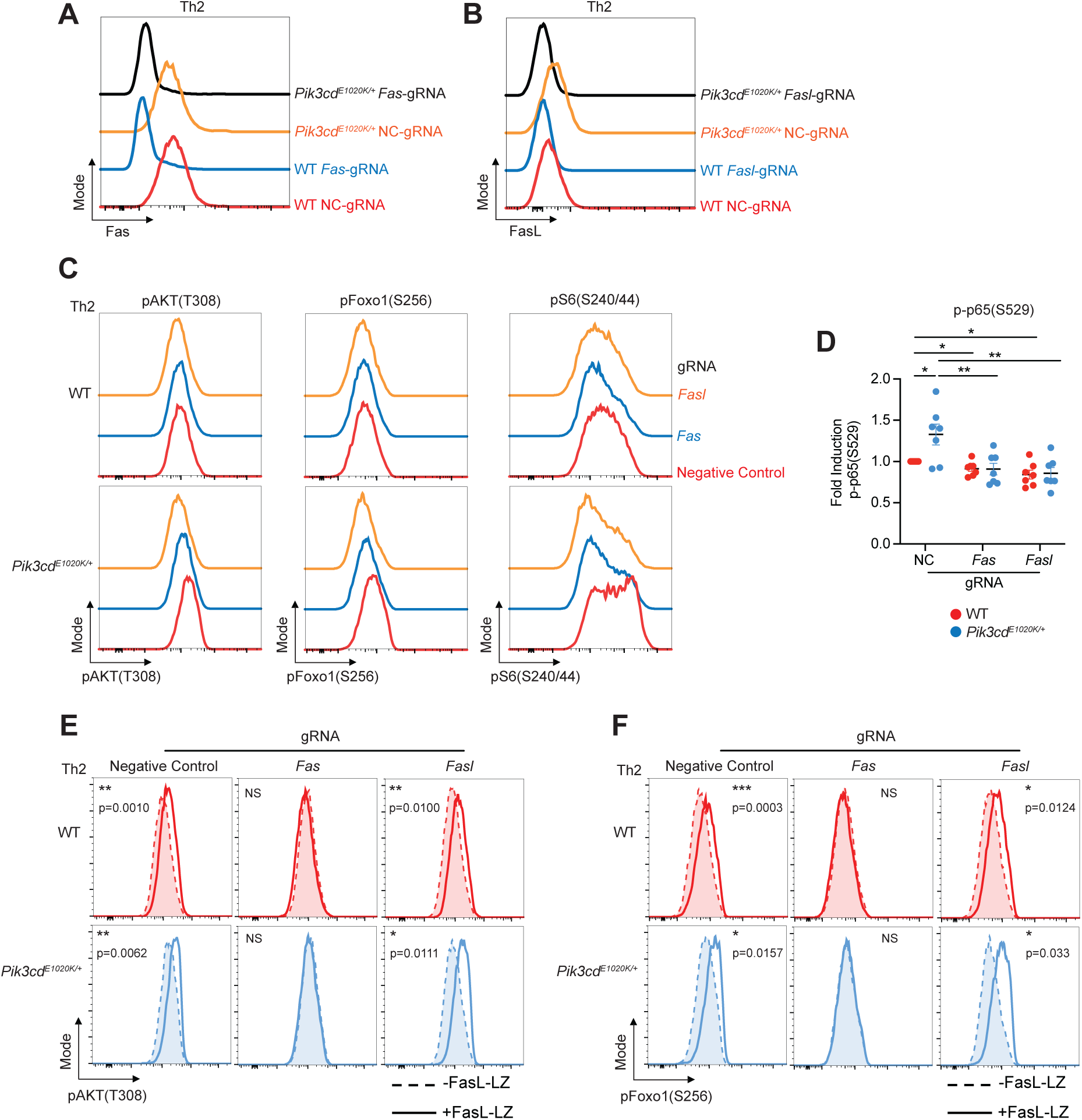
Fas-FasL signaling potentiates T cell activation. A-B) Naïve CD4 T cells from WT and *Pik3cd^E1020K/+^* mice were nucleofected with Cas9-gRNA complexes containing negative control, *Fas* or *Fasl* targeting gRNAs and underwent Th2 polarization. A) Representative flow cytometry histograms showing surface Fas expression in the indicated groups. B) Representative flow cytometry histograms showing surface FasL expression in the indicated groups. C) Representative flow cytometry histograms showing pAKT(T308), pFoxo1(S256) and pS6(S240/44) staining in negative control, *Fas* and *Fasl* gRNA targeted Th2 polarized live CD4^+^ T cells from the indicated mice. Supporting data for Figure 6F. D) Fold induction of p-p65(S529) in Th2 polarized liveCD4 T cells from the indicated groups, measured by flow cytometry. For all readouts, fold induction was calculated by normalizing MFIs to negative control WT cells. n=7 for each group, 7 independent experiments. E-F) Dashed lines represent cells cultured without FasL-LZ, solid lines show cells cultured with FasL-LZ. Supporting data for Figure 6H. E) Representative flow cytometry histograms showing pAKT(T308) staining in Th2 polarized live CD4^+^ T cells from the indicated groups. F) Representative flow cytometry histograms showing pFoxo1(S256) staining in Th2 polarized liveCD4 T cells from the indicated groups. Statistical comparisons were made using ratio paired t-tests. *p<0.05, **p<0.01, ***p<0.001, ****p<0.0001.

**Supplemental Figure 7.**
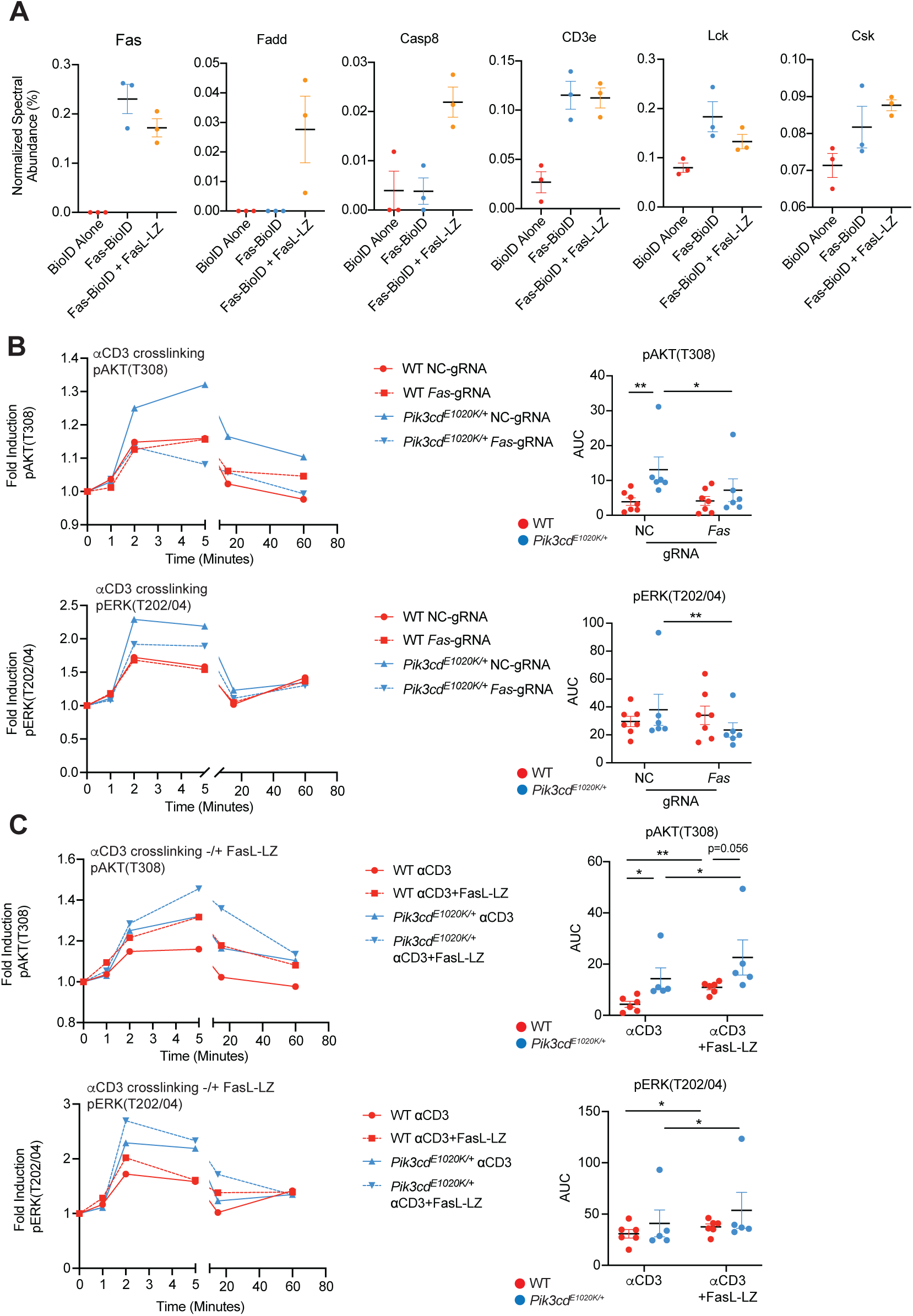
Fas interacts with the TCR complex and co-stimulates TCR signaling. A) Normalized spectral abundance (%) of the indicated proteins in BioID alone, Fas-BioID and Fas-BioID + FasL-LZ Jurkat cells, measured by mass spectrometry. B) Naïve CD4^+^ T cells from WT and *Pik3cd^E1020K/+^* mice were nucleofected with Cas9-gRNA complexes containing negative control or *Fas* targeting gRNAs and underwent aCD3/CD28 stimulation in the presence of hIL-2 for 72 hours. Cells were subsequently rested in serum free media, underwent aCD3 (1μg/mL) crosslinking over a time course (0, 1, 2, 5, 15, 60 minutes) and were analyzed by flow cytometry. n=6-7 for each group, 6-7 independent experiments. Left, fold induction of pAKT(T308) and pERK(T202/04) over time for the indicated groups. Fold induction was calculated using MFIs normalized to the 0 timepoint of the corresponding sample. Right, area under the curve (AUC) quantification of pAKT(T308) and pERK(T202/04) time courses for the indicated groups. C) Naïve CD4^+^ T cells from WT and *Pik3cd^E1020K/+^* mice underwent aCD3/CD28 stimulation in the presence of hIL-2 for 72 hours. Cells were subsequently rested in serum free media, underwent aCD3 (1μg/mL) crosslinking in the presence or absence of FasL-LZ over a time course (0, 1, 2, 5, 15, 60 minutes) and were analyzed by flow cytometry. n=6-7 for each group, 6-7 independent experiments. Left, fold induction of pAKT(T308) and pERK(T202/04) over time for the indicated groups. Fold induction was calculated using MFIs normalized to the 0 timepoint of the corresponding sample. Right, AUC quantification of pAKT(T308) and pERK(T202/04) time courses for the indicated groups. Statistical comparisons were made using ratio paired t-tests. *p<0.05, **p<0.01.

